# Multiscale integration of environmental stimuli in plant tropism produces complex behaviors

**DOI:** 10.1101/2020.07.30.228973

**Authors:** Derek E. Moulton, Hadrien Oliveri, Alain Goriely

**Affiliations:** Mathematical Institute, University of Oxford, Oxford OX2 6GG, United Kingdom

**Keywords:** Plant tropism, Biomechanics, Morphoelasticity, Rod theory, Mathematical model

## Abstract

Plant tropism refers to the directed movement of an organ or organism in response to external stimuli. Typically, these stimuli induce hormone transport that triggers cell growth or deformation. In turn, these local cellular changes create mechanical forces on the plant tissue that are balanced by an overall deformation of the organ, hence changing its orientation with respect to the stimuli. This complex feedback mechanism takes place in a three-dimensional growing plant with varying stimuli depending on the environment. We model this multiscale process in filamentary organs for an arbitrary stimulus by linking explicitly hormone transport to local tissue deformation leading to the generation of mechanical forces and the deformation of the organ in three dimensions. We show, as examples, that the gravitropic, phototropic, nutational, and thigmotropic dynamic responses can be easily captured by this framework. Further, the integration of evolving stimuli and/or multiple contradictory stimuli can lead to complex behavior such as sun following, canopy escape, and plant twining.

**Significance Statement:** To survive and to thrive, plants rely on their ability to sense multiple environmental signals, such as gravity or light, and respond to them by growing and changing their shape. To do so, the signals must be transduced down to the cellular level to create the physical deformations leading to shape changes. We propose a multiscale theory of tropism that takes multiple stimuli and transforms them into auxin transport that drives tissue-level growth and remodeling, thus modifying the plant shape and position with respect to the stimuli. This feedback loop can be dynamically updated to understand the response to individual stimuli or the complex behavior generated by multiple stimuli such as canopy escape or pole wrapping for climbing plants.

---

Plant tropism is the general phenomenon of directed growth and deformation in response to stimuli. It includes phototropism, a reaction to light (1); gravitropism, the reaction to gravity (2, 3); and thigmotropism, a response to contact (4), among many others (see Fig. 1). The study of tropisms in plants dates back to the pioneering work of giants such as Darwin (5) and Sachs (6), and has been a central topic for our understanding of plant physiology ever since. Tropisms form a cornerstone subject of modern plant biomechanics (7), crop management strategies (8), as well as systems biology and plant genomics (9). Being sessile by nature, plants lack the option to migrate and must adapt to their ever-changing environment. The growth response of individual plants to environmental cues will determine the yield of a crop in unusually windy conditions, will decide the future of rainforests in a world driven by climate change, and may be key for colonizing foreign environments such as Mars.

**Fig. 1.**
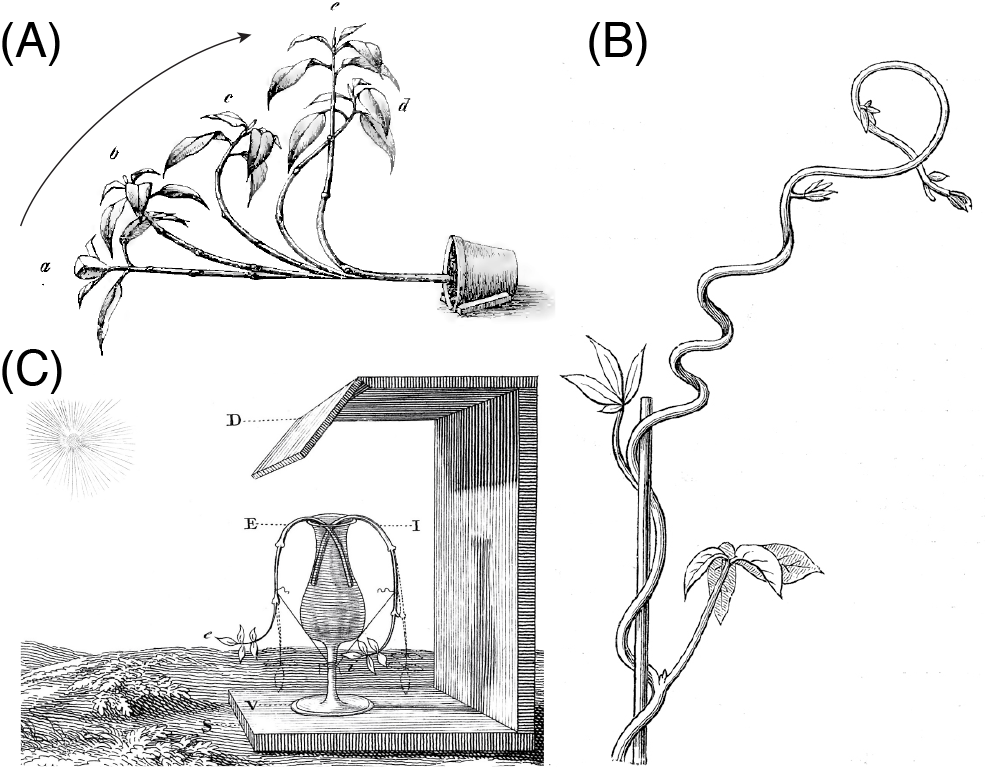
Classic experiments on tropic responses. (A) Gravitropism: a potted plant realigns itself with gravity (13). (B) Thigmotropism: a twining vine develops curvature when in contact with a pole (6). (C) Phototropism: a plant reorients itself towards the light source (18th Century experiments by Bonnet (14), correctly interpreted by Duhamel du Monceau (15, 16)).

Mathematical modeling plays an invaluable role in gaining a better understanding of tropisms and how plants may respond to a change in their environment (10). Yet, a general mathematical description of tropisms is a grand challenge. First, the growth response tends to be dynamically varying: a sunflower grows to face the sun, but as it grows the sun moves, so the environmental influence – the intensity of light impacting on each side of the sunflower – is changing during the process. Similarly, a tree branch may align with the vertical in a gravitropic response; decreasing the likelihood of breaking under self-weight; however, the growth response itself may increase the branch weight and thus change the stimulus (11). Second, while there exist numerous experimental setups that enable to carefully isolate a particular stimulus, a plant typically receives multiple stimuli at the same time and in different locations (12). The resulting movement is an integration of multiple signals and movements. Third, any tropism is fundamentally a multiscale phenomenon. Transduction of an environmental cue takes place from the organ to the cell and involves, ultimately, molecular processes. A hormonal response is induced, which leads to different cells expanding at different rates in response to the chemical and molecular signals. However, one cannot understand the change in shape of the plant and its position in relation to the direction of the environmental stimulus at this level. To assess the effectiveness of the growth response, one needs to zoom out. The net effect of a non-uniform cell expansion due to hormone signaling is a tissue-level differential growth (1) as depicted in Fig. 2. At the tissue level, each cross section of the plant can be viewed as a continuum of material that undergoes non-uniform growth and/or remodeling (17). Differential growth locally creates curvature and torsion, but it also generates residual stress (18). In this way the global shape of the plant, as well as its material properties, evolve. To characterize this global change, and to update the position of the plant in the external field, a further zooming out to the plant or organ level is appropriate. At the plant level, the global shape, material properties, and positioning in the external stimulus are well described by a physical filament: here, the plant is viewed as a space curve endowed with physical properties dictated by the lower level tissue scale, and its shape and motion can be described by the theory of elastic rods, that has been applied to multiple biological contexts, from DNA and proteins, to physiology and morphogenesis (19–22).

**Fig. 2.**
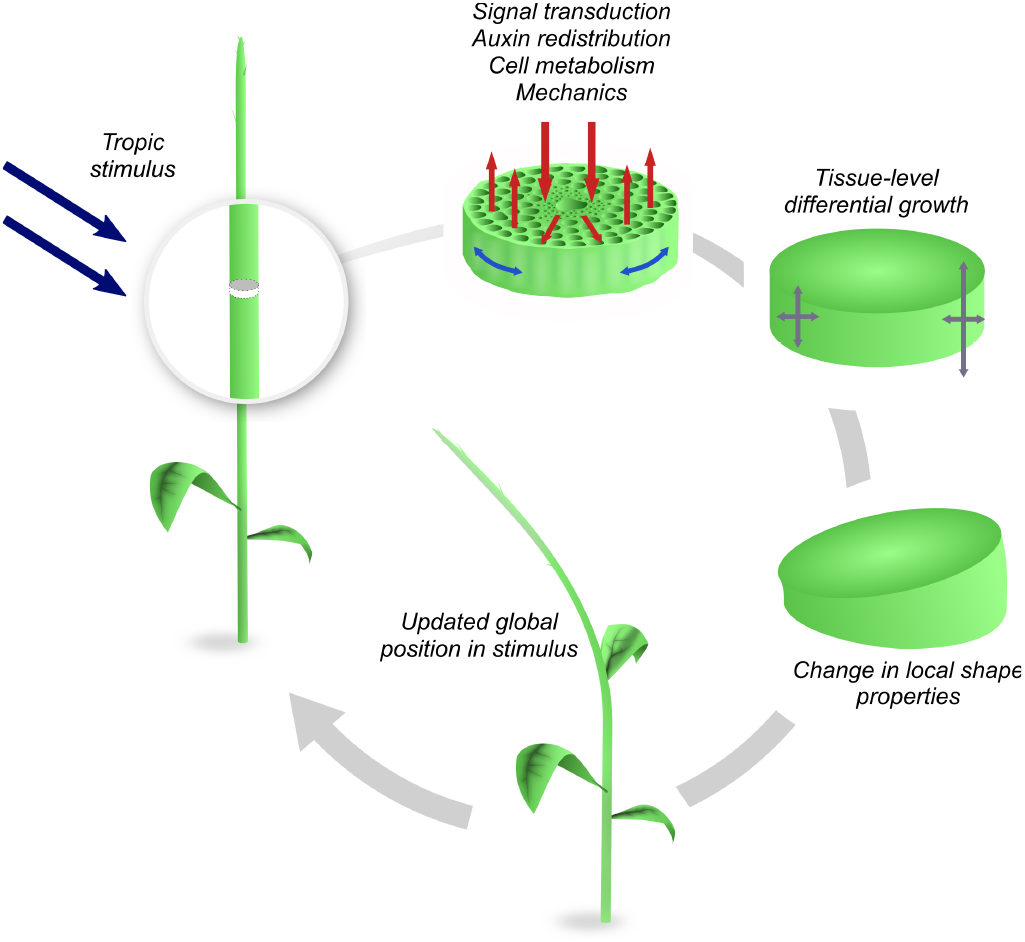
Tropism is a multiscale dynamic process: the stimulus takes place at the plant or organ level and its information is transduced down to the cellular level creating a tissue response through shape inducing mechanical forces that change the shape of the organ. In the process, the plant reorients itself and, accordingly, the stimulus changes dynamically.

The challenge of formulating a mathematical model of tropism is further complicated by the extraordinary variation in plants and the multiple types of tropism. Within a single plant, a tropic response may refer to the growth and movement of the entire plant, or a subset: a single branch, vine, stem, or root. Here, we use the word ‘plant’ to refer to the entire class of plant structures that may undergo such growth responses. Moreover, even within a single plant, multiple environmental cues will combine and overlap in effecting mechanotransductive signals, hormonal response, differential growth, and ultimate change in shape (23); e.g. a sunflower exhibiting phototropism still perceives a gravitational signal.

At the theoretical level, a variety of approaches have recently been proposed. Growth kinematics models successfully describe the tropic response at the plant-level (7, 24, 25), but do not include mechanics and cellular activities. A number of elastic rod descriptions of tropic plant growth have also been proposed (26–31); these involve a full mechanical description at the plant-level, with phenomenological laws for the dynamic updating of intrinsic properties such as bending stiffness and curvature, but specific cell- and tissue-level mechanisms are not included. Multiscale formulations have also appeared, including Functional-Structural Plant Models (8, 32, 33) and hybrid models with vertex-based cell descriptions (34, 35). These computational approaches have the potential to incorporate effects across scales but are limited to small deformations compared to the ones observed in nature.

The goal of this paper is to provide a robust mathematical theory that links scales and can easily be adapted to simulate and analyze a large number of overlapping tropisms for a spectrum of plant types. Our mathematical and computational framework includes (*i*) large deformations with changes of curvature and torsion in three-dimensional space; (*ii*) internal and external mechanical effects such as internal stresses, self-weight, and contact; and (*iii*) tissue-level transport of growth hormone driven by environmental signals. By considering the integration of multiple conflicting signals, we also provide a view of a plant as a problem-solving control system that is actively responding to its environment.

## 1. Multiscale modeling framework

The key to our multiscale approach is to join three different scales: stimulus-driven auxin transport at the cellular level; tissue-level growth mechanics; and organ-level rod mechanics.

### A. Geometric description of the plant

We start at the organ scale and model the plant as a growing, inextensible, unshearable elastic rod. This is a one-dimensional filamentary object that can bend and twist with some penalty energy, but also extend without stretching by addition of mass. Let **r**(*S, t*) ∈ R^3^ describe its centerline, where *S* is the initial arc length measured from the base of the plant towards its tip (see Fig. 3(A)). Along this centerline, we define an orthonormal basis {**d**_*i*_}, *i* = 1, 2, 3, oriented such that **d**_3_ aligns with the tangent *∂***r**/*∂S* in the direction of increasing *S*, and (**d**_1_, **d**_2_) denote material directions fixed within each cross-section. From the director basis, the Darboux vector is defined as **u**= u_1_**d**_1_ + u_2_**d**_2_ + u_3_**d**_3_, and encodes the rod’s curvature, torsion and twist (17). For a given curvature vector, the shape of the rod is determined by integrating the system of equations

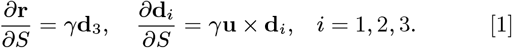

**Fig. 3.**
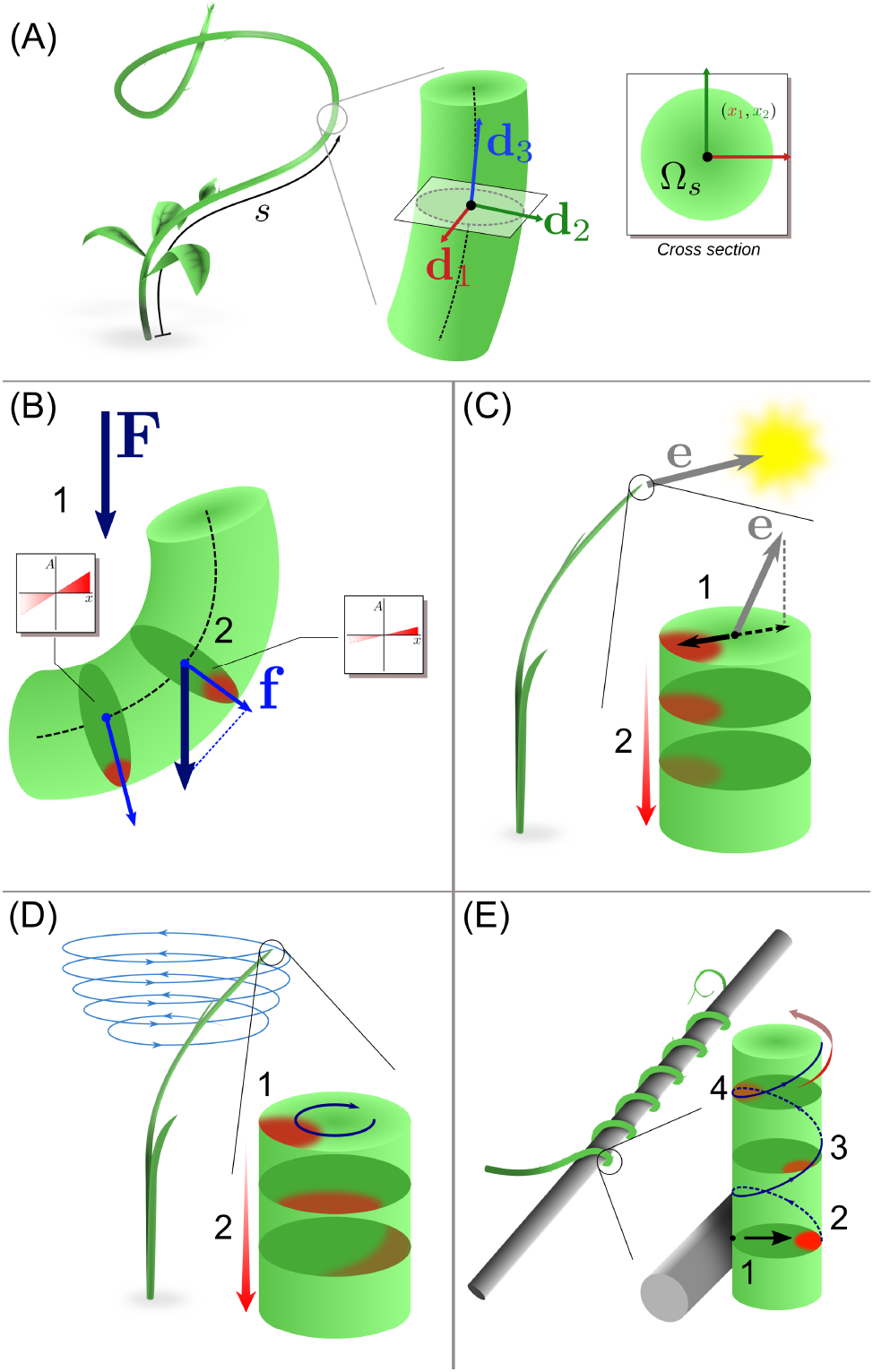
(A) Each cross section Ω*s* of the rod is parametrized with its arc length *s* (oriented acropetally) and equipped with a local material basis {**d**_1_, **d**_2_, **d**_3_}. (B) Gravitropism: the gravity vector (1) is sensed in each cross section and causes lateral auxin flow (2). (C) Phototropism: the light vector is sensed at the plant apex and results in the establishment of an apical auxin profile (1) that is transported basipetally with attenuation (2). (D) The circumnutation is generated by an internal oscillator with pulsation *ω* associated with rotating auxin profile at the apex (1). The apical profile is transported basipetally (2), generating curvature and torsion. (E) Thigmotropic pole wrapping is triggered by a contact (1) eliciting an asymmetrical auxin profile (2), which is in turn transported helically (3) to the rest of the plant with signal attenuation (4).

Here *γ* ≔ *∂s/∂S* denotes the total axial growth stretch of each section mapping the initial arc length *S* to the current arc length *s* (36). At each value of *S*, the cross section is defined by a region (*x*_1_, *x*_2_) ∈ Ω_*S*_ ⊂ *R*^2^, where *x*_1_, *x*_2_ are local variables describing the location of material points in the respective directions **d**_1_, **d**_2_. In terms of the local geometry, any material point **X** = (*X*_1_, *X*_2_, *X*_3_) in the plant can be represented by its arc length *S* and its position (*x*_1_, *x*_2_) on the cross section at *S* as follows:

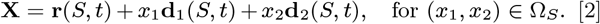

We can now use this representation to formulate the stimuli.

### B. The stimuli

Tropic stimuli are characterized by their origin, sign, and direction (37). Signal *origin* includes: chemicals, water, humidity, gravity, temperature, magnetic fields, light, touch. Tropisms can have a *sign*: *positive* if the plant grows towards or in the direction of the stimulus or *negative* if it moves away from the stimulus. The *direction* of tropism describes the orientation of the response with respect to a directed stimuli: *exotropism* is the continuation of motion in the previously established direction, *orthotropism* is the motion in the same line of action as the stimulus, and *plagiotropism*, is the motion at an angle to a line of stimulus.

Physically, stimuli are fields acting in space at a point **X** = *X*_1_**e**_1_ + *X*_2_**e**_2_ + *X*_3_**e**_3_ ∈ R^3^ and changing over time *t*. They can be either scalar fields, *f* = *f* (**X**, *t*), e.g. chemical, temperature, or light intensity; vector fields, **F** = **F**(**X**, *t*), e.g. geomagnetic field, gravity, or light direction; and, possibly, tensor fields (e.g. mechanical stress–not considered here). These stimuli are in general functions of both space and time which makes plant tropism a physical theory of fields (which is appropriate since plants grow in physical fields). Since the stimulus is defined at points in space, we must take into account the orientation and the position of the plant in space. For example, the cellular response to light in phototropism is linked to the relative orientation of the plant in relation to the light source. In the case of a vector stimulus **F**, we must therefore decompose the stimulus in the local basis:

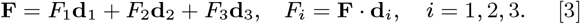

Next, we link an external stimulus to the cellular response.

### C. Cellular response and auxin transport

At the cellular level, deformation takes place through an anelastic expansion of the cell walls in response to turgor-induced tension (38, 39). We refer to any geometric change of cellular shape as growth. While detailed models of these cellular processes are available (40–42), we adopt here a coarse-grained view in which the anelastic expansions are connected *locally* via a single hormone concentration field that plays the role of a morphogen (for simplicity, we use the phytohormone *auxin*, but any other hormone could be used, or, more realistically, a combination of hormones, interacting with heterogeneous gene expression patterns). Indeed, laterally asymmetrical auxin redistribution is broadly accepted as a universal mechanism underlying tropisms (43, 44). A lateral gradient is controlled via the relocalization of auxin transporters in response to tropic signals (45). In shoots, higher levels of auxin are generally associated with faster growth. The resulting asymmetrical growth of cells elicits global curvature at the organism level through pathways that are not completely understood (46). Therefore, in our model auxin flux is a function of tropic signal and growth is taken to depend only on auxin concentration.

We model the effective migration, synthesis and uptake of auxin through a standard reaction-advection-diffusion equation (47) for the auxin concentration *A*(*x*_1_, *x*_2_, *S, t*):

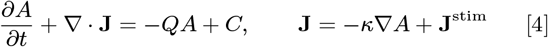

where **J** is the auxin flux, *Q* is the uptake of auxin, and *C* captures any sources or sinks. The flux is a sum of a *diffusive component* **J**^diff^ = −*κ* ∇ *A*, and a *stimulus component* **J**^stim^. Depending on the particular tropism, the information about the stimuli is contained either in **J**^stim^, or in a boundary or source term. The auxin transport equation (Eq. 4) is combined with a no-flux condition **J · n** = 0 at the outer boundary of each cross-section, where **n** is an outward normal vector to the boundary *∂*Ω_*S*_.

### D. Tissue-level growth and remodeling

Once the auxin distribution is known from the solution of Eq. (4), we can relate the growth field at the tissue level to the concentration *A*. Here, we use the general theory of morphoelasticity (17) that assigns at each point of the plant a growth tensor dictating the deformation due to growth. Physically, this tensor field integrates the multiple contributions of local pressure, cell material properties and tissue geometry, all regulated via the cell metabolic and genetic activity into a single object describing the local change of shape of an elementary volume element (48). This growth tensor may be different in different directions (anisotropic growth) and/or spatially varying (heterogeneous growth) (49). Since we are interested in change of curvature and torsion, we assume that growth only takes place, locally, along the axial direction and not in the cross-sectional direction (hence prohibiting a change in thickness). In that case, the only non-trivial component of the growth tensor is a single function *g*(*x*_1_, *x*_2_, *S, t*) that describes the change of axial length of an infinitesimal volume element (see SI Section 1). An initially straight filament of length *L*_0_ with *g* constant at all points would grow to a new straight filament of length *L* = *L*_0_(1 + *g*) (18). If, however, *g* varies from point to point, the same filament would tend to bend and twist as shown in Fig. 2.

Next, we connect the axial growth function *g* to the concentration of auxin *A* via a growth law of the form

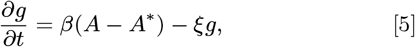

where *A*\* is a baseline level of auxin, and *β* characterizes the rate at which an increase in auxin generates growth. The extra term −*ξg* models *autotropism*, the tendency to grow straight. Indeed, this term introduces a decay of the signal in time so that the total growth stretch is a function of a time-integrated auxin signal

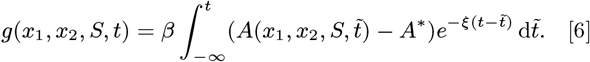

### E. Change in local shape and properties

The axial growth function *g* is defined at the tissue scale and, as such, does not directly give the change in curvature and torsion of the plant. Indeed, the change of shape depends not only on *g* but also on the internal mechanical balance of the forces generated by each growing volume element. Following the general theory given in (18) and its adaptation to the particular case of plants given by the growth law Eq. (5), we compute the intrinsic curvatures and elongation of the growing plant (SI Section 2). In the absence of autotropism (*ξ* = 0), these curvatures (given by the vector 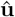) define the shape of the plant in the absence of body force and external loads:

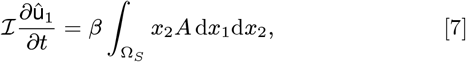

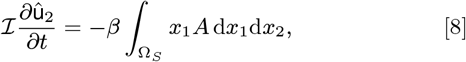

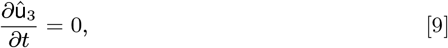

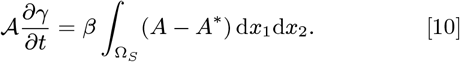

Here and for the rest of the paper, we have assumed that the cross section is circular with radius *R*, area 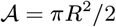 and second moment of area 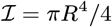.

### F. Rod mechanics sets the plant position in the stimulus field

Once the intrinsic curvatures and elongation of the plant following growth have been updated, the plant position and orientation are updated by solving the Kirchhoff equations (36, 50) for the balance of linear and angular momentum for given external forces such as self-weight or wind, and subject to any kinematic or contact constraints (see SI Section 3 for details).

Since at this scale the plant is treated as a one-dimensional structure, its equilibrium shape is easily computed even in complex non-planar geometries. Once the deformation is determined, the multiscale cycle is completed by updating the map between external stimulus and cell-scale response with respect to the updated orientation, and the process repeats.

### G. Summary

The multiscale flow of information for a given stimulus field proceeds as follows:

I. given an initial plant shape, a stimulus **F** impacts auxin transport and thus local concentrations of auxin via the transport equation (4);
II. the local auxin concentration *A* changes the local growth field that impacts the intrinsic curvatures and axial extension of the plant via Eqs. (7)–(10);
III. the new intrinsic curvatures and external conditions determine the new mechanical equilibrium of the plant, thus changing the plant position and shape in the stimulus field.

To illustrate how complex plant morphologies and tropic responses may be efficiently simulated and analyzed within this framework, we show how this multiscale process can be applied to a diversity of tropic responses and combined stimuli.

## 2. Examples

### A. Gravitropism

Gravitropism has been extensively studied both experimentally and theoretically. The classic description is called the ‘sine rule’ in which the change in curvature follows the sine of the angle with the direction of gravity (51). While it is successful in capturing observed behavior in gravitropic experiments, it is mostly phenomenological and is only applicable to planar geometry. Here, we show that the sine rule emerges naturally from our formulation but that it can be generalized to include three-dimensional deformations that are generated when a plant is rotated in time.

The stimulus for negative gravitropism is the vector field **F** = −*G***e**_3_ which can be written in the plant frame of reference as **F**= **f**+f_3_**d**_3_ where **f**≔ f_1_**d**_1_ + f_2_**d**_2_ is the gravity force acting in the plane of the cross-section. As it has been proposed that plants are insensitive to the strength of gravitational field, it is sufficient to use a unit vector representing only the direction of gravity, i.e. we scale the gravitational acceleration *G* to 1. If **f ≡ 0**, no tropic response will occur. Gravity perception relies on specific cells called *statocytes* distributed along the shoot (52). Statocytes contain dense organelles (*statoliths*) that sediment under the effect of gravity. Tilting of the plant causes statoliths to avalanche and to form a free surface perpendicular to the gravity vector, providing orientational information to the cell (53). It has been observed that the gravitropic response depends upon the angle between the statoliths free surface and the vertical, but not on the intensity of the gravitational field and the pressure of statoliths against the cell membrane (54). A possible mechanism is that the contact between the statoliths and the cell membrane may trigger relocalization of PIN membrane transporters and a redirection of auxin flux (53). Here, we follow this hypothesis and, accordingly, assume that gravity drives an advective flow of auxin **J**^stim^ = *kA***f**. If the statocytes are uniformly distributed within the stem volume then *k* is constant. We assume also a source and sink of auxin at each cross-section, representing a continual axial auxin flow, and that auxin transport is advection-dominated with timescales much shorter than the one associated with growth. Combining the transport equation (4), growth law (5) in the absence of autotropism (*ξ* = 0), and the evolution laws given by Eqs. (7)–(10), we obtain (see SI Section 4) the *gravitropic curvature law* and axial extension law:

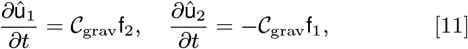

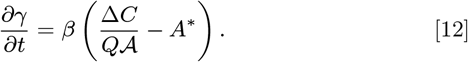

Here, 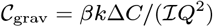 is a single constant characterizing the rate of change of curvature due to gravity and associated with a timescale of gravitropic reaction 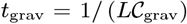, where *L* is a characteristic axial length, say the length of the plant. Note that the right hand side of Eq. (12) is proportional to the net ‘excess auxin’, the integral of (*A − A*\*) over the section, while the quantity *A*\* does not appear in Eq. (11). Thus, an excess of auxin is needed for axial growth, while only a redistribution of auxin is needed to generate curvature.

In the particular case where the plant can only bend around the single axis **d**_2_ and all external forces can be neglected, we have f_2_≡0, 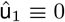 and the curvature 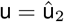. Defining *α* to be the inclination angle, Eq. (11) reads

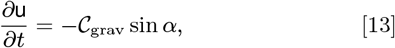

which is the classic and widely-used sine law of gravitropism (24).

The general laws Eqs. (11) and (12) can be used for more complex gravitropic scenarios. Consider, for instance, an experiment in which the base of the plant stem is at a fixed angle *θ* from the horizontal and the base is rotated, as shown in Fig. 4(A). Then, in the frame of reference of the plant, the direction of gravity is constantly changing. Here, we consider the case of zero-axial growth and neglect self-weight (see Section C for these extra effects). The tropic response will generate curvature and torsion depending on the angle and the rotational velocity of the base as shown in Fig. 4. For visualisation purposes, we fix the base rotation rate to one turn per unit time and vary the tropic reaction rate of the plant, which is equivalent to a fixed reaction rate and varying base rotation rate via a rescaling of time. In Fig. 4, we simulate three full rotations of the base with varying reaction rates (see also SI movies 1-4). The evolving morphology is characterized by three metrics: an alignment metric in (B) that measures how closely aligned with the vertical the plant is (a value of one is attained if the entire plant is vertical), and curvature and torsion in (C) that broadly measure deviation from a straight configuration (details in SI Section 5).

**Fig. 4.**
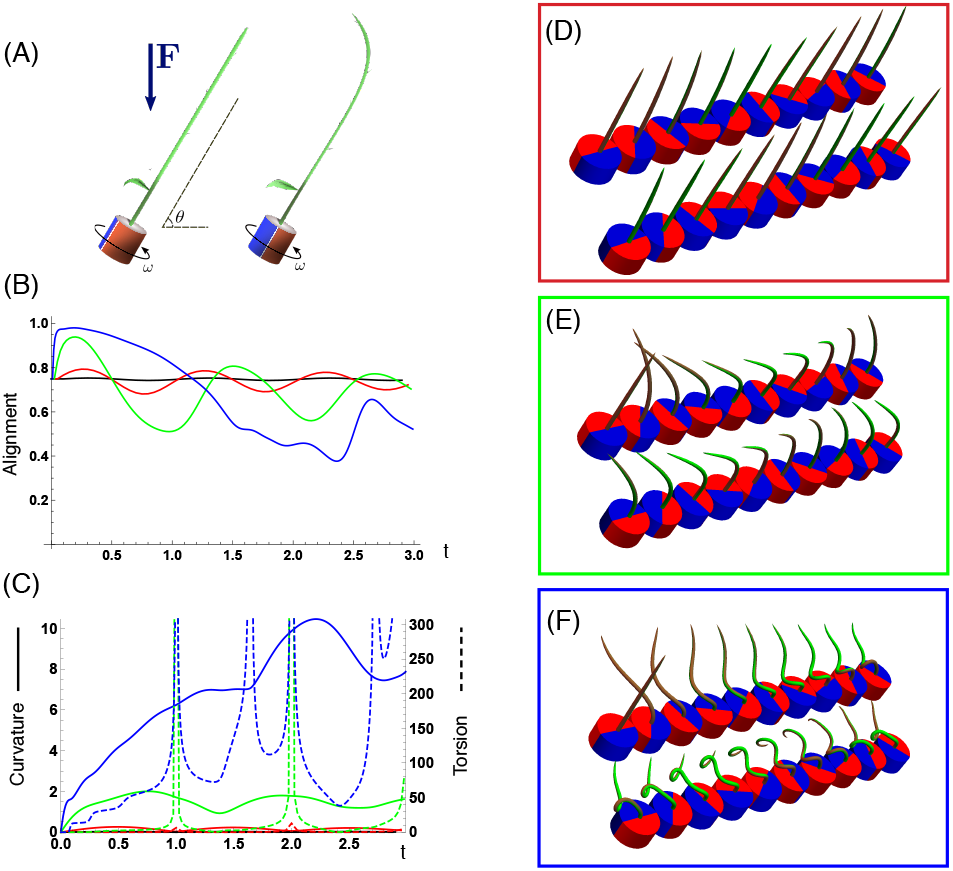
Gravitropism with a rotating base. (A) A base is tilted with respect to the vertical and then rotated about the axis with speed so that one revolution is completed every time unit. Gravitropic response is simulated for varying values of gravitropic sensitivity 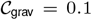 (black), 1 (red), 10 (green), and 50 (blue). Alignment with the vertical (B) and curvature and torsion (C) are plotted against time for three base rotations. Snapshots for cases of (D) slow reaction, 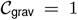, (E) intermediate reaction, 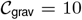, and (F) fast reaction, 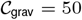. The sequence is read left to right, top to bottom, and the base rotation is counterclockwise. Further simulation details and parameters provided in SI Section 7.

Consider first the s lowest reaction morphology (equivalent to the case of fastest base rotation), given by the black curves in Fig. 4(B)-(C). Since the plant’s response time is much slower than the base rotation, the gravitropic response is averaged out and the plant hardly deviates from the straight configuration, never improving its alignment and generating effectively no curvature or torsion. The plant is almost perfectly straight at all times (snapshots not included). The red curves denote a case with increased but still small reaction (fast base), which generates only small oscillations in alignment and curvature. In this regime, the plant is effectively ‘confused’; the local gravitational field is changing too quickly for the plant to make any progress towards alignment with the gravitational field.

As the reaction rate is increased (or the base rotation decreased), interesting morphologies emerge. In the case of the intermediate reaction rate _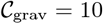_ (green curves), the plant begins to curve towards the vertical during the first quarter rotation of the base, bending about the **d**_2_ axis and increasing its alignment. However, as the base continues to rotate, the curvature initially developed has the tip pointing away from the vertical, so the alignment decreases, and the plant now must bend about the orthogonal **d**_1_ axis. As the base completes its first rotation and the ‘desired’ axis for bending returns to the original **d**_2_, an inversion occurs (more visible in the movies provided in SI), creating a large spike in torsion. This basic process repeats with each rotation.

Finally, increasing the reaction rate (or slowing the base) further creates highly complex morphologies as evidenced by the blue curves. Here the plant quickly aligns with gravity and attains near perfect alignment in the first tenth of the first rotation. As the base rotates away from this aligned state, we see an interesting phenomenon: the tip of the plant is able to react and maintain alignment with the vertical, but since the base of the plant is clamped at an ever-changing angle, a loop forms starting at the base and working its way to the tip. This is accompanied by strong variations in the total alignment and increasingly high curvature, with repeated spikes in torsion as extra twist is removed. Our simulations of this case beyond three rotations suggest that while the basic process of loops generated at the base and working to the tip continues, the morphology does not settle down into a fixed oscillatory pattern, highlighting the potential for complex dynamics generated by this highly nonlinear system.

### B. Phototropism

It was Darwin, at the end of the 19th Century, who demonstrated that exposure of the plant apex to a light source was necessary to induce tropic bending (5, 55). Later on, Peter Boysen-Jensen proposed that bending is induced by a diffusive substance, later identified as auxin, that carries the tropic information from the apex to the rest of the shoot (56, 57). These early observations are the basis of the popular Cholodny-Went model (58, 59) stating that phototropism relies upon three broad mechanisms: *(i)* sensing of light direction at the tip of the shoot; *(ii)* establishment of a lateral asymmetry of auxin concentration at the tip; and *(iii)* basipetal transport of this asymmetrical distribution, resulting in differential growth along the shoot (60–62).

We model these three steps by considering *axial* transport of auxin, with an asymmetrical distribution that is established at the shoot apex by the stimulus, treated as a point source of light. We suppose that auxin flows basipetally with advective velocity *U* and uptake *Q*, and assuming slow diffusion, the transport equation is

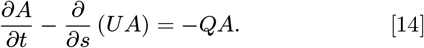

where the derivative in space is taken with respect to the current arc length *s* ∈ [0, *ℓ*]. Additional source/sink terms can be used to model axial extension without changing the evolution of the curvature but are omitted in the first instance. We account for the amount and distribution of auxin at each section via a boundary condition at the tip (*s* = *ℓ*) and define *A*_tip_(*x*_1_, *x*_2_, *t*) = *A*(*x*_1_, *x*_2_, *ℓ, t*) that depends on the light source located at **p**(*t*) in space, and a scalar *I*(*t*) representing its intensity. We then define the unit vector **e** from the plant tip to the light source and write it in the plant reference frame: **e**(*t*) = e_1_**d**_1_(*ℓ*)+ e_2_**d**_2_(*ℓ*)+ e_3_**d**_3_(*ℓ*), as shown in Fig. 3(C). The line e_1_*x*_1_ + e_2_*x*_2_ in the cross section distinguishes the light side of the tip from the dark side and defines the asymmetrical distribution of auxin:

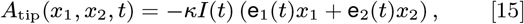

where *κ* characterizes the sensitivity of the phototropic response.

For constant velocity *U*, and in the absence of autotropic effects, Eqs. (14) and (15)) can be solved exactly (SI Section 4), which gives the *phototropic curvature law*:

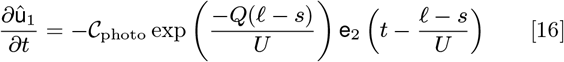

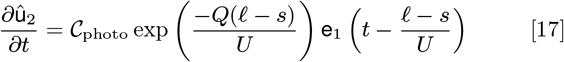

where 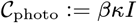 is a single parameter characterizing the rate of curvature, and from which the phototropic response time is defined as 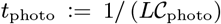. The exponential decay in Eq. (16) is due to the uptake of auxin so that less is available at the base, while the time shift *t*_tran_ ≔ *L/U* of *A*_tip_ accounts for the transport time to the section at arc length *s*, leading to time-delay differential equations. The change of curvature thus depends on three quantities: *(i)* the orientation of the tip with respect to the light source *t* = (*ℓ − s*)/*U* ago; *(ii)* the amount of auxin available for the phototropic signal, which depends on the uptake *Q*; and *(iii)* the plant’s response sensitivity, characterized by 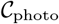. Bending occurs over a characteristic dimensionless bending length *ℓ*_bend_ ≔ *U/Ql* within the tip of the plant.

#### B.1. Fixed light source–no growth

We consider first a fixed light source and restrict our attention to the case of zero axial growth so that *ℓ* = *L* for all time and the transport equation is solved in the reference variables. For a given transport time *t*_tran_, the response of the plant is determined by the characteristic bending length *ℓ*_bend_ and the response time as shown in Fig. 5. With small bending length, the response is localized close to the tip, and the plant is much slower to orient (comparing (A) and (B)). Increasing the response rate naturally produces a faster orientation and potentially an overshoot. If axial auxin transport is much faster than the growth response (*t*_tran_ ≪ *t*_photo_), then the auxin is effectively in steady state at each growth step (as we have assumed for the cross-sectional transport). This implies that the delay can be neglected and the curvature response at each point depends on the *current* orientation of the tip. In this case, since the response is characterized entirely by the orientation of a single point, the motion is very simple: the plant bends to orient with the light, with no oscillations about the state in which the tip is perfectly oriented with the light (**e**_2_ = 0).

**Fig. 5.**
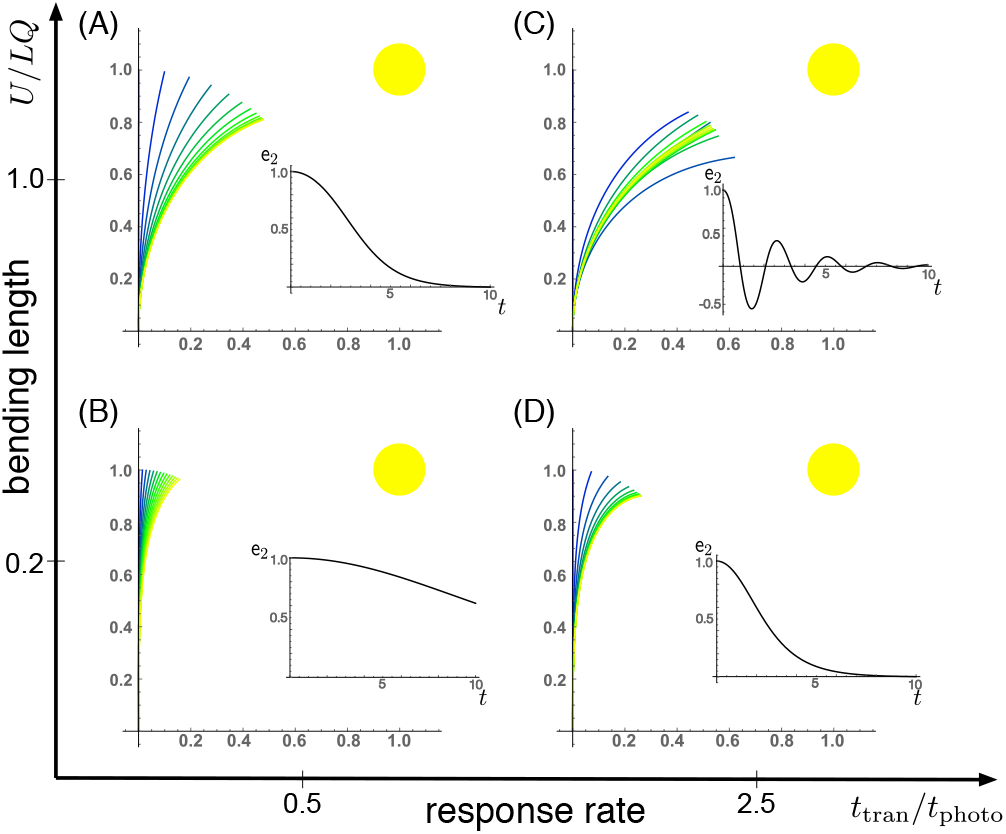
Planar phototropic shape evolution for a fixed light source and with no axial growth, for small and large values of the ratio of response rate to transport rate *t*_tran_ /*t*_photo_ and dimensionless bending length *ℓ*_bend_ = *U/QL*. Inset: alignment is characterized by **e**_2_ (*t*), such that **e**_2_ = 0 when the tip is pointed at the source for this planar case. Further simulation details and parameters provided in SI Section 7.

Contrast this behavior with gravitropism, in which an oscillation about the vertical state is typical unless a strong autotropism response is added. The difference between the gravitropic law and the phototropic law for fast transport is that during gravitropism, each cross section tries to align *itself* with gravity, thus creating a conflict at the global level that results in an oscillatory motion; while during phototropism each cross section tries to align the tip with the light, so there is no conflict. However, with delay, such a conflict does exist, due to the fact that each cross section is accessing a previous state of the tip. Thus, in the regime *t*_tran_ ~ *t*_photo_, and if the bending length is not too short, a damped oscillation about the preferred orientation is observed, as shown in Fig. 5(C).

#### B.2. Moving light source

Next, we consider a moving source, and in particular we simulate a day-night cycle of a plant following a light source (the sun) as shown in Fig. 6(A). The intensity *I*(*t*) is also taken to be sinusoidal, so that the phototropic signal is strongest at noon and the signal vanishes at sunset. For fast response and long bending length, the plant bends significantly and succesfully tracks the moving light source (Fig. 6(B)). However, at night and without a signal the motion halts (see also SI movie 5). The plant remains bent towards sunset the entire night and does not display the nocturnal reorientation observed in many plants (63, 64). With a non-vanishing autotropic term *ξ* in Eq. (5), we obtain an *autophototropism curvature law* of the form

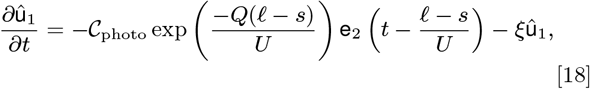

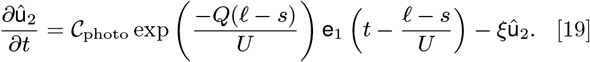

**Fig. 6.**
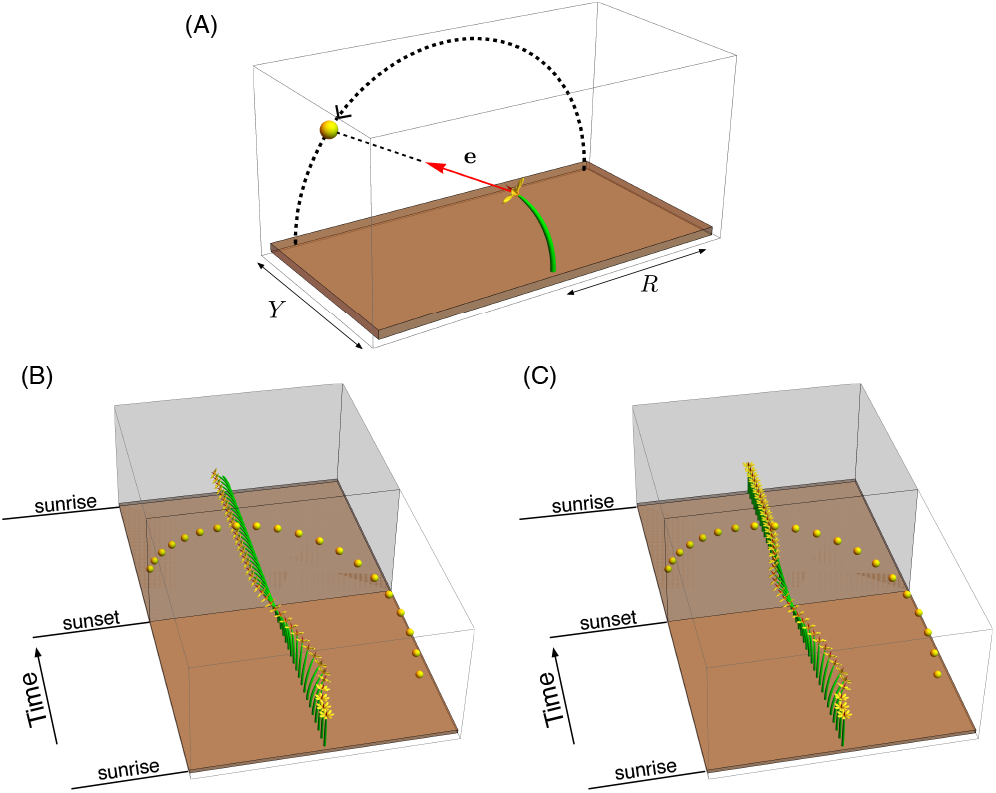
(A) Geometry of the phototropic response stimulated by a lightsource that follows a circular path of radius *R*, shifted a distance *Y* in the transverse ‘horizon’. (B) A full day-night cycle with fast response and long bending length. (C) the addition of autotropic terms enables the plant to return to the vertical during night, when the phototropic signal is absent. Further simulation details and parameters provided in SI Section 7.

The additional terms serve to straighten the plant in the absence of any other signal. This is evident in Fig. 6(C), in which we see that the motion during the day is very similar, while at night the stem straightens back to the vertical (see also SI movie 6). We note that in heliotropic plants such as the common sunflower, *Helianthus annuus*, there are additional mechanisms, not considered here, based on circadian rhythms to reorient the plant at night to face eastward in anticipation of the next sunrise (65).

### C. Photogravitropic response

Next, we demonstrate the delicate balance that must exist in the presence of tropic responses to multiple stimuli. We simulate two different scenarios of a plant responding to simultaneous but conflicting gravitropic and phototropic signals. Following (12), we assume that the effects of multiple stimuli are additive (see SI Section 4E). This assumption is based on the existence of separate pathways for signal transduction leading to the redistribution of auxin. However, it is known that these pathways share common molecular processes and there are non-trivial interactions between different tropisms (66) that will not be included here.

#### C.1. Fixed horizontal light source

We consider a growing plant subject to self-weight and initially oriented vertically, but with a fixed light source located in the transverse horizontal direction. The evolution of the plant can then be characterized by the ratio of response rates to gravitropic versus phototropic signals, and the ratio of density to bending stifness, which controls the degree of deformation under self-weight.

In Fig. 7(A)-(D) we show the evolving morphology of the plant in this two-dimensional parameter space, plotting both the deformed shape (solid lines) and the reference unstressed shape (dashed lines). In (A), (B), the effect of self-weight is relatively minimal, and the evolution is primarily driven by the conflicting phototropic signal acting horizontally to the right, and the vertical gravitropic signal. With increased mass, there is an increased mechanical deformation due to self-weight, so that significant disparity develops between the deformed and reference shapes. In this regime the balance of signals has greater importance for the fate of the plant. Comparing (B) and (D), the initial phases are similar, but as the plant lengthens and extends to the right in (D), self-weight deforms the plant significantly, with half of the plant below the base level by the end of the simulation. Such a deformation could signal failure by creating large torque at the base. In (E) we plot the moment at the base, where the stress is highest, against time for each case, and as expected the moment is significantly higher for larger mass.

**Fig. 7.**
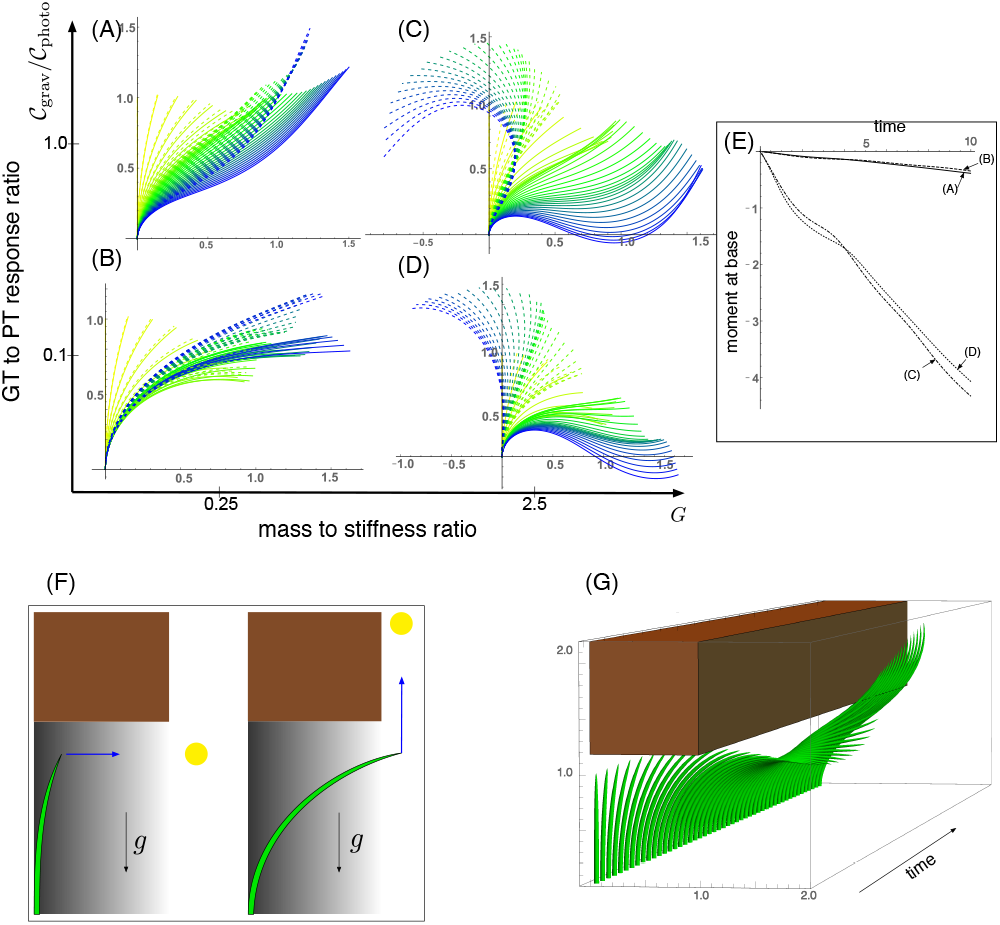
Gravitropism versus phototropism. In (A)-(E), a fixed light source is located to the right, at the point (4,1), while gravity points vertically downward. For each parameter set, the tropic response is simulated for the same total time and with equivalent axial growth. The evolving plant shape, deformed under self-weight, is shown in increasing time from yellow to blue, with the unstressed shape appearing as a dashed line. The moment at the base for the 4 cases is plotted against time in (E). (F) depicts the setup for a plant escaping from the shade under a rigid obstacle. The phototropic signal either points horizontally, if the tip is under the shade, or vertically, if the tip is out of the shaded region. (G) a sample simulation showing a successful escape. Further simulation details and parameters provided in SI Section 7.

Intuitively, we expect that this problem could be alleviated by increasing the gravitropic response rate. Comparing (C) and (D), the evolution with higher gravitropic response in (C) does show decreased sagging. However, the moment at the base is in fact higher in (C). Increasing the gravitropic response rate even further does ultimately alleviate the problem – consider that the plant remains mostly vertical if gravitropism dominates phototropism – nevertheless, this example highlights the delicate and potentially counter-intuitive nature of this balance.

#### C.2. Escaping from the shade

The results above suggest a view of a plant as a problem-solving agent that actively responds to the signals in its environment. A typical problem that many plants have to solve is access to light. For instance, consider a plant growing underneath a canopy as shown in Fig. 7(F) and (G). While the tip is in the shaded region, diffuse lighting creates a phototropic stimulus to grow horizontally, orthogonal to the gravitational signal. If the tip emerges from under the shade, phototropism and gravitropism align, and the plant will attempt to grow vertically. In this mixed signal scenario, the success or failure of the plant in emerging from under the canopy is down to how the competing signals are integrated, and the relative importance of self-weight. An example of a successful escape is shown in Fig. 7(G). Note that determining the mechanical forces acting in the plant is crucial in this example: once the tip is outside of the shade, both signals try to align the entire length of the plant with the vertical, and this leads to physical contact between the plant and the corner of the canopy. Determining the morphology beyond this point thus requires determining the mechanical contact force (see SI Section 6) that would not be possible in a purely kinematic description.

### D. Pole dancing

A fascinating plant motion is the mesmerizing dance that climbing plants, such as twiners, perform to first find a pole and then wrap around it. Like any dance, this event requires a complex integration of stimuli to achieve a well-orchestrated sequence of steps: *(i)* finding a pole, *(ii)* contacting the pole, and *(iii)* proceeding to wrap around the pole. A common mechanism for searching for a climbing frame is circumnutation, a combination of circular movement and axial growth causing the tip to move up in a sweeping spiral path, as first described by Darwin (5, 26, 67). When the plant makes contact with a pole, it must then interpret its orientation with respect to the pole in order to wrap around it. Here the stimulus is mechanical: the physical contact of the plant with the pole results in a change of curvature, a response referred to as thigmotropism. Since initially the plant only samples a very small region of the pole, the stimulus field is highly localized. As the plant wraps around the pole, new contact points are established to propagate the helical shape upward.

#### D.1. Circumnutation

Experiments suggest that, depending on the plant, circumnutation is either driven by an internal oscillator, a time-delay response to gravity, or a combination of the two (68, 69). Here, following the hypothesis of an internal oscillator, we show that the basic nutating motion emerges naturally from an internal oscillator at a single point combined with axial auxin transport (70). We consider an auxin source at the point *s* = *s_c_* from which an auxin differential is transported axially. The auxin transport equation may be solved in a similar manner as in the phototropism case with an added rotational component in the local frame of the cross section due to an internal oscillator (Fig 3(D)). Taking for simplicity a constant rotation rate *ω*, we obtain (details in SI Section 4) the *circumnutation curvature law*:

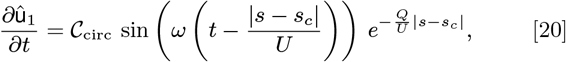

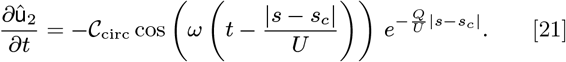

Since the signal here is internal, there is no feedback from the environment and the morphology of the plant is predetermined by the uptake *Q*, the transport velocity *U*, the response rate 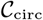, and the rotation rate *ω*. In Fig. 8(A) we illustrate a sample motion with auxin source at the tip. Fig. 8(B) demonstrates the impact of auxin uptake: high uptake means the motion is constrained to a region very close to the tip and thus the elliptical shape of the tip pattern is small. More complex tip patterns may also be generated if there is nonuniformity in the internal oscillator (Fig. 8(C)).

**Fig. 8.**
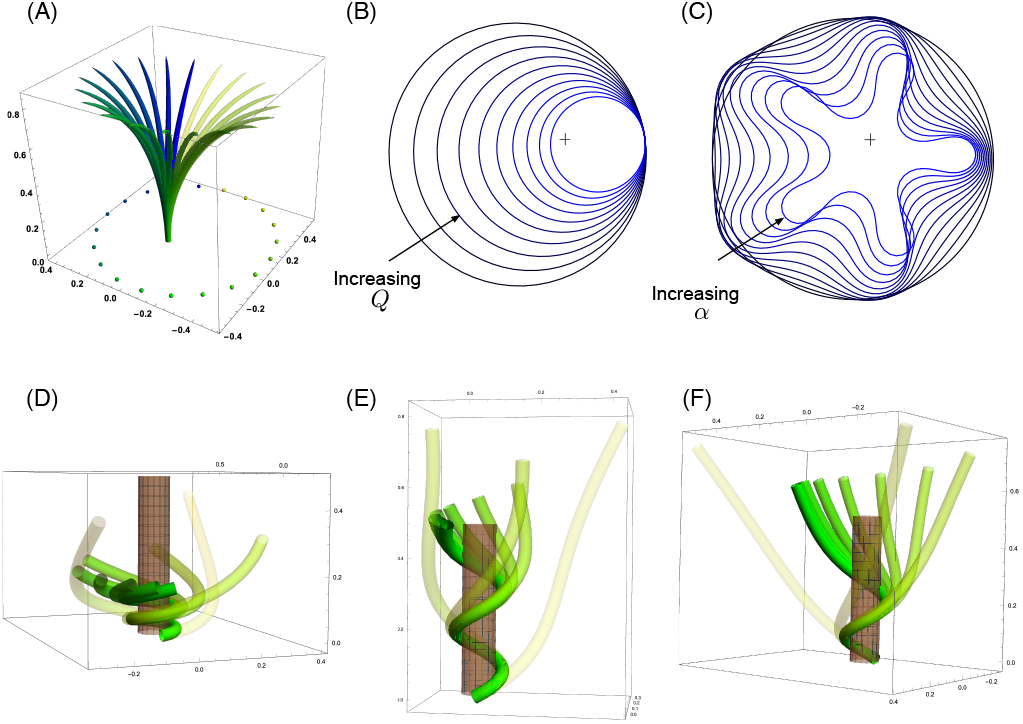
Circumnutation (A)-(C) and thigmotropism (D)-(F). (A) snapshots of a sample circumnutation motion, with tip pattern projected onto the plane. In (B) and (C), tip patterns are plotted for varying parameters (the location of the plant base is indicated by the cross). In (B), an increase in uptake *Q* decreases the size of the tip pattern. In (C), a non-constant angular velocity of the oscillator is given by 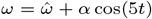, generating a tip pattern with 5-fold symmetry, and increasingly non-circular with increasing *α*. In (D)-(F), the wrapping around a pole due to thigmotropism via a single contact point is simulated for the same total time, for different parameter regimes: with low rotational component (D), the torsion is low, a high rotational component with low uptake (E) generates rapid wrapping and high torsion, while wrapping is much slower with high uptake (F). Further simulation details and parameters provided in SI Section 7.

#### D.2. Thigmotropism

Two interesting observations can be made when a twining plant makes first contact with a pole: *(i)* torsion is generated via a localized contact around a single point, and *(ii)* a rotation is induced, i.e. the orientation of the tangent of the plant with respect to the axis of the pole changes. These observations suggest that this contact is sufficient to generate locally a helical shape, and that the pitch of the helix is fixed by internal parameters as opposed to the angle at which contact is made (71, 72).

To show how pole wrapping can be obtained within our framework consistently with these observations, we consider a plant with a single contact point located at *s* = 0, and at position on the boundary Ω_0_ with angle *ψ*_0_ in the plane **d**_1_−**d**_2_. The contact induces an auxin gradient at this point, with maximal auxin on the opposite side of the contact point, i.e. *A*(0, *x*_1_, *x*_2_) = −*F* (cos *ψ*_0_ *x*_1_ + sin *ψ*_0_ *x*_2_), and the auxin is transported by an advective flux with both an axial component *U* and a constant rotational component with angular velocity *ω* (Fig 3(E); see also SI Section 4). The transport equation can be solved exactly (SI Section 4), and we obtain the following *thigmotropism curvature law*:

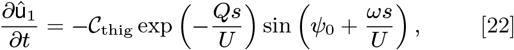

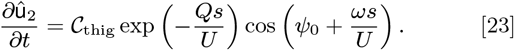

Solving these equations leads to exact expressions for the intrinsic curvatures from which we extract the geometric curvature 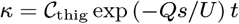 and torsion *τ* = *ω/U* (see SI Section 4). The curvature increases linearly in time until it reaches a maximal value determined by the pole radius (the intrinsic curvature may keep increasing, but the actual curvature may not due to the mechanical contact). For a pole of radius *c* and plant radius *a* the helix radius *α* = *c* + *a* is fixed, while the helical angle *ϕ* depends on the rotation rate and is found to satisfy sin(2*ϕ*) = *ωα/U*.

For given axial velocity *U*, the resulting helical shape is determined solely by the geometry of the pole and the rotational component *ω*, while the wrapping rate depends on the uptake *Q* and the response rate 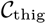. In Fig. 8(D)-(F) we illustrate three different regimes: low rotational component with low uptake (D), high rotational component with low uptake (E), and high rotational component with high uptake (F) (see also SI movies 7-9). Note that at time 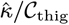 where 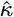 is the final curvature, the contact point spreads to a contact region, creating a wave of contact and auxin signal that propagates along the length of the plant. Here, we restrict our attention to the signal from the first contact point. The separate curvature laws for circumnutation and pole wrapping can now be combined to simulate the process of searching for a pole, making contact, and wrapping (see SI movie 10).

## 3. Conclusion

Plant motion in response to environmental stimuli is a process of extreme biological and ecological relevance. While the pioneering biologists of the 19th Century investigated the global motion of plants via clever experiments devised to create conflicting signals and generate complex plant morphologies, most of the work of the 20th Century was focused on the molecular and cellular processes, seeking signaling pathways and relevant proteins involved in tropic response. We have combined this accumulated knowledge with recent progress in the physical and computational modeling of living structures to develop a general theory of tropism that relates stimuli to shape. To do so, we created a framework that combines information about auxin transport and the mechanisms by which environmental stimuli are integrated into cellular activities, tissue-level growth, and ultimately a change in shape at the plant scale represented by a morphoelastic structure.

We have demonstrated the power of this framework through a series of examples including key effects such as axial growth, autotropism, gravitropism, phototropism, thigmotropism, self-weight, circumnutation, contact mechanics, and threedimensional deformations. The specific tropic scenarios we have considered were chosen to illustrate the range of complex behaviors capable of being simulated. The study of individual stimuli provided new laws for the evolution of curvatures for different form of tropisms. Further, we demonstrated the potential for multiple and potentially conflicting stimuli to create *problems* for the plant to *solve*. The resulting plant behaviors indicate a need for a delicate balance between competing tropisms to achieve a particular task. In each particular scenario, we have opted for parsimony over complexity in terms of modeling choices, in order to highlight qualitative features and how auxin-level differences may become apparent in plant-level morphology. Nevertheless, our approach is fully compatible with more detailed modeling choices, though potentially at the cost of increased computation time.

The work presented here integrated information at the tissue and organ levels. This approach needs to be expanded to include cell-based and molecular-level descriptions to fully link the scales for tropisms and plant growth. At the cell to tissue scale, the link between auxin and growth in Eq. (5) is a lumped description of a complex process that involves cell wall tension and turgor pressure (73). In principle, additional modeling layers could also be added between the stimulus and auxin response, including for example transcription factors and protein production and interactions. Both of these extensions likely require explicit cell-based modeling. However, our framework is such that if the output from a cell model is the value of the piecewise continuous axial growth function *g*, then the evolution of the curvatures given by Eqs. (7)–(9) still applies and can be used to infer the global changes of geometry.

This work provides a theoretical platform for understanding plant tropisms and generating complex morphologies. As well as linking to cellular and subcellular scales, a key future direction is connecting with experiments dedicated to controlling multiple stimuli and generating complex morphologies. For instance, we showed that a relatively simple experimental set-up like a rotating base under gravity can generate a wide range of plant shapes. Such steps espouse an approach that is both multiscale and multidisciplinary. Indeed, plants refuse to obey by the rules of a single scientific discipline. They are not simply genetic or cellular entities, nor are they purely physical objects or ecological atoms. They reach for the sun, they bend under gravity, they feel their neighbors, they grow, twist, curve, and dance in the fresh air and in the dark caves. If we have any hope to understand them, we will need to respect their plurality, break down our own disciplinary barriers, and fully integrate our scientific knowledge from subcellar to ecological levels.

## ACKNOWLEDGMENTS

The support for A.G. by the Engineering and Physical Sciences Research Council of Great Britain under research grant EP/R020205/1 is gratefully acknowledged. The authors thank János Karsai for permission to use his flower parameterisation in Fig. 6.

