## Supplementary Information for "Multiscale integration of environmental stimuli in plant tropism produces complex behaviors"

### Supplementary Information for *Multiscale integration of environmental stimuli in plant tropism produces complex behavior*

**Derek E. Moulton, Hadrien Oliveri, and Alain Goriely**

Mathematical Institute, University of Oxford, Oxford OX2 6GG, United Kingdom

#### 1. From 3D growth field to 1D elasticity

In obtaining the curvature evolution laws, we follow the framework of (1) to map from a growth tensor on a 3D tubular structure to the intrinsic curvature and axial growth of the same object when viewed as a 1D elastic rod. In this framework, note that we define the centerline along the centroid of each cross section so that

$$\int_{\Omega_S} x_1 \, \mathrm{d}x_1 \mathrm{d}x_2 = \int_{\Omega_S} x_2 \, \mathrm{d}x_1 \mathrm{d}x_2 = \int_{\Omega_S} x_1 x_2 \, \mathrm{d}x_1 \mathrm{d}x_2 = 0. \tag{1}$$

We consider a growth tensor of the form

$$\mathbf{G} = G_{ij} \mathbf{e}_i \otimes \mathbf{e}_j, \quad i, j = 1, 2, 3,$$

where in general each $G_{ij}$ may be functions of position, and $(\mathbf{e}_1, \mathbf{e}_2, \mathbf{e}_3)$ are Cartesian basis vectors that are chosen to coincide with the frame $(\mathbf{d}_1, \mathbf{d}_2, \mathbf{d}_3)$ in the initial pre-deformed state of the plant.

Conceptually, the components of $\mathbf{G}$ define the expansion (or contraction) of material both as a local property and as directional quantities. For example, if $G_{11} > 1$ then cells will expand in the $\mathbf{d}_1$ direction. The determinant of $\mathbf{G}$ describes the net gain or loss of mass at each point; here it is worthwhile to note that growth without a change of mass is possible, and it is also possible to have a growth field for which points gain or lose mass while the total mass for a cross section stays fixed.

The quantity that is of most relevance for capturing a tropic growth response is the term $G_{33}$; this term describes axial elongation; it is heterogeneity of this term across a section that generates curvature in the plant. While other terms may play a role, for instance in changing the cross-sectional geometry, we posit that this will typically be a secondary effect, and thus to make progress we consider the growth tensor to take the form*

$$\mathbf{G} = \mathrm{diag}(1, 1, 1 + g). \tag{2}$$

For the growth tensor defined above, the key result of (1) is that the intrinsic curvatures are given explicitly by

$$\mathcal{I}_1 \hat{\mathsf{u}}_1(S, t) = \int_{\Omega_S} x_2 g(x_1, x_2, S, t) \, \mathrm{d}x_1 \mathrm{d}x_2, \quad \mathcal{I}_2 \hat{\mathsf{u}}_2(S, t) = -\int_{\Omega_S} x_1 g(x_1, x_2, S, t) \, \mathrm{d}x_1 \mathrm{d}x_2, \quad \hat{\mathsf{u}}_3 = 0, \tag{3}$$

where

$$\mathcal{I}_1 := \int_{\Omega_S} x_2^2 \, \mathrm{d}x_1 \mathrm{d}x_2, \quad \mathcal{I}_2 := \int_{\Omega_S} x_1^2 \, \mathrm{d}x_1 \mathrm{d}x_2$$

are the second moments of area. A straightforward extension of the derivation given in (1) shows that the axial elongation $\gamma$ is given by

$$\mathcal{A} \gamma = \int_{\Omega_S} g(x_1, x_2, S, t) \, \mathrm{d}x_1 \mathrm{d}x_2, \tag{4}$$

where $\mathcal{A}$ is the cross-sectional area. Note in particular that if $g$ is constant, then from Eq. (1) it follows that no curvature is generated, and the axial extension is equal to $g$; this reflects the simple and intuitive notion that uniform axial growth does not create bending.

#### 2. Obtaining curvature and growth evolution laws

As described in the main text, the axial growth $g$ is connected to auxin concentration $A(x_1, x_2, s, t)$ by a growth law, assumed to have the form

$$\frac{\partial g}{\partial t} = \beta (A - A^*). \tag{5}$$

*In many cases the framework is still compatible with non-zero growth in terms other than $G_{33}$, though the solution method will be greatly complicated if these induce residual stress, see (1).

Since the shape of the cross sections is assumed to remain constant in time, we can take a time derivative across equations Eqs. (3) and (4) and utilize Eq. (5) to obtain evolution laws for the curvatures:

$$\mathcal{I}_1 \frac{\partial \hat{\mathsf{u}}_1}{\partial t} = \beta \int_{\Omega_S} x_2 A \,\mathrm{d}x_1 \mathrm{d}x_2, \;\; \mathcal{I}_2 \frac{\partial \hat{\mathsf{u}}_2}{\partial t} = -\beta \int_{\Omega_S} x_1 A \,\mathrm{d}x_1 \mathrm{d}x_2, \;\; \frac{\partial \hat{\mathsf{u}}_3}{\partial t} = 0. \tag{6}$$

And similarly, the evolution law for the axial extension is

$$\mathcal{A} \frac{\partial \gamma}{\partial t} = \beta \int_{\Omega_S} (A - A^*) \,\mathrm{d}x_1 \mathrm{d}x_2. \tag{7}$$

The approach outlined in (1) allows for more generic growth tensor $\mathbf{G}$, in which case the bending stiffnesses about the $\mathbf{d}_1$ and $\mathbf{d}_2$ axes, as well as the torsional stiffness, can also change due to the growth. However, for the growth form Eq. (2), the stiffnesses are not impacted by $g$. Hence, for this modeling choice, when passing from tissue to organ scale, the tropic response is entirely encoded by the change in the intrinsic curvature of the plant as well as any axial extension.

An extension of this model to include *autotropism*, which consists in adding a decay term

$$\frac{\partial g}{\partial t} = \beta (A - A^*) - \xi g. \tag{8}$$

This extra term $-\xi g$ models the tendency to grow straight in the absence of other signals and the total growth stretch is then a function of a time-integrated auxin signal

$$g(x_1, x_2, S, t) = \beta \int_{-\infty}^{t} (A(x_1, x_2, S, \tilde{t}) - A^*) e^{-\xi(t - \tilde{t})} \,\mathrm{d}\tilde{t}. \tag{9}$$

#### 3. Rod geometry and Kirchhoff equations

The output of the tissue-level modeling is a set of evolution laws for the intrinsic curvatures $\hat{\mathsf{u}}_i$ and the axial growth stretch $\gamma$. The actual morphology of the plant is then determined by solving the Kirchhoff equations for an elastic rod with non-zero evolving intrinsic curvature and axial growth. Here we briefly recall the basic elements of rod theory. A *rod* is a space curve $\mathbf{r}(S) \in \mathbb{R}^3$, known as the *centerline*, equipped with two additional unit orthonormal vector fields $(\mathbf{d}_1(S), \mathbf{d}_2(S))$ representing the orientation of a cross section at $S$. The *general frame* is obtained by defining $\mathbf{d}_3(S) = \mathbf{d}_1(S) \times \mathbf{d}_2(S)$ and we note that $\{\mathbf{d_1}, \mathbf{d_2}, \mathbf{d_3}\}$ forms a right-handed orthonormal basis. The components of a vector $\mathbf{a} = \mathsf{a}_1 \mathbf{d}_1 + \mathsf{a}_2 \mathbf{d}_2 + \mathsf{a}_3 \mathbf{d}_3$ in the local basis are denoted by $\mathsf{\mathbf{a}} = (\mathsf{a}_1, \mathsf{a}_2, \mathsf{a}_3)$. We note that $|\mathbf{a}| = |\mathsf{\mathbf{a}}|$.

We choose the material parameter $s$ to be the current arc length, i.e. in the grown configuration, and $S$ to be the material arc length in an initial pre-grown configuration. These are related by the growth stretch

$$\gamma := \frac{\partial s}{\partial S}. \tag{10}$$

For an unshearable rod, we may choose $\mathbf{d}_3$ to align with the tangent direction, so that

$$\frac{\partial \mathbf{r}}{\partial s} = \mathbf{d}_3, \tag{11}$$

or equivalently

$$\frac{\partial \mathbf{r}}{\partial S} = \gamma \mathbf{d}_3. \tag{12}$$

A complete kinematic description of the frame is given by:

$$\frac{\partial \mathbf{d}_i}{\partial s} = \mathbf{u} \times \mathbf{d}_i, \quad i = 1, 2, 3, \tag{13}$$

where $\mathbf{u}$ is the *Darboux vector*. The first two components $(\mathsf{u}_1, \mathsf{u}_2)$ of the Darboux vector are associated with the Frenet curvature while $\mathsf{u}_3$ represents twisting, that is the rotation of the basis (not the curve) around the $\mathbf{d}_3$ vector. It contains both information on the Frenet torsion $\tau$ of the centerline and on the rotation of the cross section for increasing values of $s$.

In particular, if the rod is assumed to be inextensible, the Darboux vector is related to the usual notion of Frenet curvature and torsion $\kappa$ and $\tau$ by

$$\cot \varphi = \frac{\mathsf{u}_2}{\mathsf{u}_1}, \tag{14}$$

$$\kappa = \sqrt{\mathsf{u}_1^2 + \mathsf{u}_2^2}, \tag{15}$$

$$\tau = \mathsf{u}_3 + \frac{\mathsf{u}_2' \mathsf{u}_1 - \mathsf{u}_1' \mathsf{u}_2}{\mathsf{u}_1^2 + \mathsf{u}_2^2}. \tag{16}$$

where prime denotes differentiation with respect to current arc length $s$. Also, $\varphi$ is the angle between the normal and the vector $\mathbf{d}_1$. The quantity $\partial\varphi/\partial s$, *the excess twist*, represents the rotation of the local basis with respect to the Frenet frame as the arc length increases.

The stress on the cross section at $\mathbf{r}(s)$ from adjacent segments with larger material coordinates ($s' > s$) gives rise to a *resultant force* $\mathbf{n}(s)$ and *resultant couple* $\mathbf{m}(s)$. These satisfy the balance of linear and angular momentum, which in mechanical equilibrium read

$$\frac{\partial \mathbf{n}}{\partial s} + \mathbf{f} = \mathbf{0}, \qquad [17]$$

$$\frac{\partial \mathbf{m}}{\partial s} + \frac{\partial \mathbf{r}}{\partial s} \times \mathbf{n} = \mathbf{0}. \qquad [18]$$

Here $\mathbf{f}$ is a linear force density accounting for any external forces acting on the rod.

The system is closed by boundary conditions and constitutive laws. We restrict to an inextensible rod in this paper, and thus only a constitutive equation relating moment $\mathbf{m}$ to curvature is needed. For a quadratic elastic energy, this takes the general form $\mathbf{m} = \mathbf{K}(\mathbf{u} - \hat{\mathbf{u}})$, where $\mathbf{K}$ is a stiffness matrix. Considering the simplest and most widely used case of a diagonal $\mathbf{K}$, we have

$$\mathbf{m} = K_1(\mathsf{u}_1 - \hat{\mathsf{u}}_1)\mathbf{d}_1 + K_2(\mathsf{u}_2 - \hat{\mathsf{u}}_2)\mathbf{d}_2 + K_3(\mathsf{u}_3 - \hat{\mathsf{u}}_3)\mathbf{d}_3. \qquad [19]$$

In this case, the Kirchhoff theory tells us that the stiffnesses are

$$K_1 = E\mathcal{I}_1, \quad K_2 = E\mathcal{I}_2, \quad K_3 = \mu J \qquad [20]$$

where $E$ is the Young's modulus, $\mu$ the second Lamé parameter and $J$, $\mathcal{I}_{1,2}$ depend on the cross-sectional shape (see main text).

In terms of boundary conditions, we primarily consider a plant that is held clamped at one end and free at the other. Denoting the clamped end $s = 0$, and the free end $s = \ell$, these amount to fixing the position and frame at $s = 0$:

$$\mathbf{r}(0,t) = \mathbf{r}_0, \quad \mathbf{d}_i(0,t) = \mathbf{d}_{i,0}, \quad i = 1,2,3, \qquad [21]$$

and imposing zero force and moment at $s = \ell$:

$$\mathbf{m}(\ell,t) = \mathbf{n}(\ell,t) = \mathbf{0}. \qquad [22]$$

As the elastic timescale is much shorter than the growth timescale, mechanical equilibrium is assumed at all times, and the intrinsic curvatures and growth stretch $\gamma$ are updated in a quasistatic fashion via a simple forward Euler time-stepping of the appropriate evolution law.

#### 4. Specific curvature evolution laws

In this section we outline the steps to obtain the curvature evolution laws given in the main text from the assumptions on auxin transport and via the general evolution equations Eqs. (6) and (7). Each tropism is considered in turn.

**A. Gravitropism.** In the case of gravitropism, we consider a gravity driven auxin flux $\mathbf{J}^{\text{stim}} = kA\mathbf{f}$, where $\mathbf{f} := \mathsf{f}_1\mathbf{d}_1 + \mathsf{f}_2\mathbf{d}_2$ describes the cross-sectional component of the direction of gravity expressed in the local frame. The parameter $k$ describes the gravitropic auxin flow rate. This is in contrast to the diffusive flux $\mathbf{J}^{\text{diff}} = -D\nabla A$, where $D$ is a diffusion coefficient. Due to the nature of our tissue-level description of auxin transport, the parameters $k$ and $D$ are difficult to quantify. At the cellular-level, models of auxin transport (2) are highly dependent on cell geometry, and auxin flux may differ significantly in the cytoplasm compared to the apoplast, due to varying diffusivity. In cell-based models of gravitropism, e.g. (3), gravitational stimulus is modeled by modifying PIN efflux carrier locations on particular cells, based on the stem orientation with respect to gravity. In this view the flux $\mathbf{J}^{\text{stim}}$ serves as a tissue-level proxy for a complex interaction of proteins and auxin transport both through and across cells; this view does not lend a natural link to the parameter $k$. Moreover, this parameter could also be related to the timescale of settling of statoliths, e.g. (4). The theoretical challenge is to bridge the vast divide that exists between cell-based descriptions of auxin transport and plant-level descriptions of tropism kinematics. The tissue-level model of auxin transport and growth mechanics we propose is a step towards bridging this divide, though, to our knowledge, tissue-level auxin transport models do not currently exist in the literature. With no available information on tissue-level parameter choices, we opt for modeling simplicity and qualitative analysis: to focus on the stimulus impact, we consider advection-dominated flow, i.e. the zero diffusion limit $D = 0$[†]. Also, as auxin transport timescales are generally shorter than the timescale associated with growth (5), and in this case transport is only occurring on the short cross-sectional lengthscale, we also take the auxin concentration to be at steady state. Under these assumptions, the auxin concentration satisfies

$$\nabla \cdot (kA\mathbf{f}) = -QA + C_{\text{in}}\delta(r - r_0) - C_{\text{out}}\delta(x_1)\delta(x_2). \qquad [23]$$

Here the divergence is only taken in the cross-sectional variables $(x_1, x_2)$, and $Q$ is the uptake. The second and third terms on the right hand side account for a source $C_{\text{in}}$ and sink $C_{\text{out}}$ of auxin in each cross section, providing a simple model of auxin transport routes. In particular, we consider here a source at radius $r = r_0$, which may for instance be taken to be near the cross-sectional radius in the case of epidermal auxin flow, and a sink at the center. These terms are needed simply to provide

[†] In the case of $D \neq 0$, the model formulation remains valid, but the techniques applied below to obtain explicit curvature evolution laws would not be. Rather, incorporating diffusion would require solving for the auxin concentration at each growth time step, which would likely require computational techniques.

a source of auxin to be transported under gravity and stimulate growth; the specifics of these choices do not impact on the resulting equations.

Combining Eqs. (5), (6) and (23), and using Eq. (1), we obtain the following equation for the intrinsic curvature $\hat{\mathsf{u}}_1$:

$$\mathcal{I}_1 \frac{\partial \hat{\mathsf{u}}_1}{\partial t} = \beta \int_{\Omega_S} x_2 (A - A^*)\, \mathrm{d}x_1 \mathrm{d}x_2 = -\frac{\beta}{Q} \int_{\Omega_S} x_2 (\nabla \cdot kA\mathbf{f})\, \mathrm{d}x_1 \mathrm{d}x_2. \quad [24]$$

Note the source and sink terms both vanish on a circular cross section, as does the $A^*$ term, assuming $A^*$ is constant. This form for $\partial \hat{\mathsf{u}}_1/\partial t$ is not very useful, as it would still require solving for the auxin concentration at each time step. However, we may determine the evolution laws without explicitly solving for $A$, by noting the following identity

$$x_2 \nabla \cdot (kA\mathbf{f}) = \nabla \cdot (x_2 kA\mathbf{f}) - \nabla x_2 \cdot kA\mathbf{f} = \nabla \cdot (x_2 kA\mathbf{f}) - kA\mathsf{f}_2, \quad [25]$$

since $\nabla x_2 = \mathbf{d}_2$. Therefore, when integrating over the cross section, we have

$$\int_{\Omega_S} x_2 (\nabla \cdot kA\mathbf{f})\, \mathrm{d}x_1 \mathrm{d}x_2 = \int_{\Omega_S} \nabla \cdot (x_2 kA\mathbf{f})\, \mathrm{d}x_1 \mathrm{d}x_2 - k\mathsf{f}_2 \int_{\Omega_S} A\, \mathrm{d}x_1 \mathrm{d}x_2 = -k\mathsf{f}_2 \int_{\Omega_S} A\, \mathrm{d}x_1 \mathrm{d}x_2, \quad [26]$$

where we have used the divergence theorem and the no-flux boundary condition $\mathbf{J} \cdot \mathbf{n} = kA\mathbf{f} \cdot \mathbf{n} = \mathbf{0}$ on $\partial\Omega_S$ to write

$$\int_{\Omega_S} \nabla \cdot (x_2 kA\mathbf{f})\, \mathrm{d}x_1 \mathrm{d}x_2 = \int_{\partial\Omega_S} x_2 kA\mathbf{f} \cdot \mathbf{n}\, \mathrm{d}s = 0. \quad [27]$$

The problem is now reduced to evaluating an integral of only $A$ over the cross section. We may again insert $A$ via Eq. (23); the divergence term again vanishes by the no-flux boundary condition, while the delta function terms integrate to a constant $\Delta C = C_{\mathrm{in}} - C_{\mathrm{out}}$, i.e. the net auxin available in the cross section, so that

$$\int_{\Omega_S} x_2 (\nabla \cdot kA\mathbf{f})\, \mathrm{d}x_1 \mathrm{d}x_2 = -\frac{k}{Q} \Delta C \mathsf{f}_2, \quad [28]$$

Combining the above, we obtain the relation provided in the main text:

$$\frac{\partial \hat{\mathsf{u}}_1}{\partial t} = \mathcal{C}_1 \mathsf{f}_2, \quad [29]$$

where $\mathcal{C}_1 = \beta k \Delta C / (\mathcal{I}_1 Q^2)$. Similar steps lead to the evolution equations for $\hat{\mathsf{u}}_2$ and $\gamma$ as appearing in the main text.

**B. Phototropism.** In the case of phototropism, as explained in the main text we consider the following axial auxin transport equation

$$\frac{\partial A}{\partial t} - \frac{\partial}{\partial s}(UA) = -QA, \quad [30]$$

and with an auxin source at the tip $s = \ell$ given by

$$A_{\mathrm{tip}}(x_1, x_2, t) = -\kappa I(t) \left(\mathsf{e}_1(t) x_1 + \mathsf{e}_2(t) x_2\right) \quad [31]$$

where $\mathbf{e}$ is a unit vector pointing from the tip to the light source, $I$ characterizes the intensity of the light, and $\kappa$ characterizes the strength of the response to generate auxin. An exact solution to Eqs. (30) and (31) is given by

$$A(x_1, x_2, s, t) = A_{\mathrm{tip}} \left(x_1, x_2, t - \frac{\ell - s}{U}\right) \exp\left(-\frac{Q(\ell - s)}{U}\right). \quad [32]$$

Following Eq. (6), we multiply by $x_2$ and integrate over a cross section. Since Eq. (31) gives $A$ as a linear function of $x_1$, $x_2$, then using Eq. (1), we obtain the curvature evolution given in the main text:

$$\frac{\partial \hat{\mathsf{u}}_1}{\partial t} = -\mathcal{C}_{\mathrm{photo}} \exp\left(-\frac{Q(\ell - s)}{U}\right) \mathsf{e}_2 \left(t - \frac{\ell - s}{U}\right), \quad [33]$$

where $\mathcal{C}_{\mathrm{photo}} = \beta\kappa I /$, and similarly for $\partial \hat{\mathsf{u}}_2/\partial t$.

In the case of phototropism with an additional autotropic term, we use the generalized growth law Eq. (9). The steps above are nearly identical, and we readily obtain the updated evolution forms

$$\frac{\partial \hat{\mathsf{u}}_1}{\partial t} = -\mathcal{C}_{\mathrm{photo}} \exp\left(-\frac{Q(\ell - s)}{U}\right) \mathsf{e}_2 \left(t - \frac{\ell - s}{U}\right) - \xi \hat{\mathsf{u}}_1, \quad [34]$$

$$\frac{\partial \hat{\mathsf{u}}_2}{\partial t} = \mathcal{C}_{\mathrm{photo}} \exp\left(-\frac{Q(\ell - s)}{U}\right) \mathsf{e}_1 \left(t - \frac{\ell - s}{U}\right) - \xi \hat{\mathsf{u}}_2. \quad [35]$$

In the formulation outlined above, there is no axial growth component, i.e. $\partial\gamma/\partial t = 0$, since the integral of $A$ over each cross-section is zero due to the form of $A_{\text{tip}}$. We may naturally incorporate axial growth by adding source and sink terms, as appeared in the gravitropism case. That is, consider the transport equation

$$\frac{\partial A}{\partial t} - \frac{\partial}{\partial s}(UA) = -QA + C_{\text{in}}\delta(r - r_0) - C_{\text{out}}\delta(x_1)\delta(x_2). \quad [36]$$

Denoting the combined source and sink terms by $\Delta C$, then if this term is independent of $s$ the solution is

$$A(x_1, x_2, s, t) = \left[A_{\text{tip}}\left(x_1, x_2, t - \frac{\ell - s}{U}\right) - \frac{\Delta C}{Q}\right]\exp\left(-\frac{Q(\ell - s)}{U}\right) + \frac{\Delta C}{Q}. \quad [37]$$

In this case, the curvature evolution laws are unchanged, while the axial growth satisfies

$$\frac{\partial\gamma}{\partial t} = \beta\left[\exp\left(-\frac{Q(\ell - s)}{U}\right)\left(\frac{\Delta C}{Q} - 1\right) - A^*\right]. \quad [38]$$

This formulation naturally produces growth focused at the tip, with growing region depending on the uptake. In cases of high uptake, it may be necessary to modify the growth law to avoid 'negative growth' ($\partial\gamma/\partial t < 0$).

**C. Circumnutation.** In the case of circumnutation, we again assume axial flow of auxin from a source point (which may naturally be the tip, but needn't be). The only difference is that the auxin gradient originating at the source has a rotational component in the cross section. In the general case, if the gradient is along the line $\cos\theta x_1 + \sin\theta x_2$, then the oscillator is described by the function $\theta = \theta(t)$. The derivation of curvature evolution is the same as in the phototropism case, simply with $\mathsf{e}_1(t)$ replaced by $\cos(\theta(t))$ and $\mathsf{e}_2(t)$ replaced by $\sin(\theta(t))$.

**D. Thigmotropism.** For thigmotropism, we assume that physical contact occurs at a point $s_c$, and with angle in the local basis $\psi_c$; that is, the point in physical space

$$\mathbf{r}(s_c, t) + R(\cos\psi_c\mathbf{d}_1(s_c, t) + \sin\psi_c\mathbf{d}_2(s_c, t)),$$

where $R$ is the cross-sectional radius. Geometrically, generating the helical shape of a twining plant requires establishing a growth gradient which rotates along the axis of the plant with increasing arc length (6). In terms of auxin transport, it has been observed that point contact creates a sharp rise in auxin concentration at the stimulus point that is transported along the stem (7). This suggests that we impose as a boundary condition at the contact point an auxin gradient, with minimum auxin at the contact point, i.e.

$$A(x_1, x_2, s_c, t) = -\kappa(\cos\psi_c x_1 + \sin\psi_c x_2), \quad [39]$$

where $\kappa$ characterizes the strength of the tropic response (which may, for instance, be connected to the magnitude of the contact force).

We then assume that auxin flux consists of a rotational cross-sectional component with angular velocity $\omega$, and an axial component with velocity $U$, thus generating a helical auxin gradient along the stem. The angular component may be seen as a proxy for (largely unknown) underlying mechanisms that generate the rotational component of growth gradient needed for helical twining. For instance, in nutating roots, a circumferential wave of ion flux is engaged; ion fluxes may interact with auxin (8), and also appear sensitive to touch (9), thus providing a possible mechanism.

Following these assumptions, the auxin transport equation is thus

$$\frac{\partial A}{\partial t} + \text{sign}(s - s_c)\frac{\partial}{\partial s}(UA) + \nabla\cdot(Ar\omega\mathbf{e}_\theta) = -QA. \quad [40]$$

Here the sign function accounts for the flow away from the contact point in either direction, the divergence $\nabla\cdot()$ is only with respect to the cross-sectional variables, $r$ is the radial position vector within a cross section, and $\mathbf{e}_\theta$ is the circumferential unit vector in the cross section. Since the curvature response is largely localized to the region near the contact point, we neglect any time delay that would occur due to axial transport, and thus consider the steady state auxin concentration. Setting $\partial A/\partial t = 0$, the exact solution is given by

$$A = -\kappa\exp\left(-\frac{Q|s - s_c|}{U}\right)\left(\cos\left(\psi_c + \frac{\omega}{U}\text{sign}(s - s_c)\right)x_1 + \sin\left(\psi_c + \frac{\omega}{U}\text{sign}(s - s_c)\right)x_2\right). \quad [41]$$

From here, the evolution equations for $\hat{\mathsf{u}}_i$ follow naturally from Eq. (6), by multiplying by $x_i$ and integrating over a cross section, again with the use of Eq. (1).

**E. Multiple signals.** We model multiple simultaneous signals as an additive effect to the growth response. In particular, consider two stimuli A and B. Letting the auxin concentration under stimulus A be denoted $A_A$, and similarly $A_B$ for stimulus $B$, the axial growth law is adapted to

$$\frac{\partial g}{\partial t} = \beta(A_A + A_B - A^*). \quad [42]$$

An alternative approach would be to formulate a single transport equation with combined flux and/or boundary conditions for each stimulus; however, this would generally necessitate fully computational techniques. The assumption of separated auxin flows for each stimulus, as utilized here, reflects the differing signal transduction pathways that exist for different stimuli, and leads to an additive growth response, as has been observed to hold reasonably well in the case of photogravitropism (10). In this way, if tropisms A and B lead to the individual curvature laws

$$\frac{\partial \hat{\mathsf{u}}_i}{\partial t} = f_A^{(i)}, \quad \frac{\partial \hat{\mathsf{u}}_i}{\partial t} = f_B^{(i)}, \ i = 1, 2 \quad [43]$$

respectively, then the curvature evolution under the combined influence of signals A and B is simply

$$\frac{\partial \hat{\mathsf{u}}_i}{\partial t} = f_A^{(i)} + f_B^{(i)}. \quad [44]$$

#### 5. Gravitropism metrics

The metrics used in quantifying the gravitropic response with a rotating base are defined as follows:

1. **Alignment** $= \frac{1}{L}\int_0^L (\mathbf{d}_3(S)\cdot\mathbf{e}_z)^2 \, \mathrm{d}S$, with $\mathbf{e}_z = (0,0,1)$
2. **Curvature** $= \frac{1}{L}\int_0^L \sqrt{\mathsf{u}_1^2(S) + \mathsf{u}_2^2(S)} \, \mathrm{d}S$
3. **Torsion** $= \frac{1}{L}\int_0^L \left(\frac{\mathsf{u}_2'(S)\mathsf{u}_1(S) - \mathsf{u}_1'(S)\mathsf{u}_2(S)}{\mathsf{u}_1^2(S)+\mathsf{u}_2^2(S)}\right)^2 \, \mathrm{d}S$

The formulas for curvature and torsion follow from Section 3.

#### 6. Escape from the shade - mechanical contact

In simulating the escape from the shade in photogravitropism (main text Fig. 7), contact with the rigid, shade-creating obstacle becomes an issue. Contact at a point $s = s_c$ induces a contact force $\mathbf{f}_c$ that must be accounted for. We assume that the plant may slide along the surface without friction, so that the contact force acts only in the normal direction. Working in a planar geometry with tangent $\mathbf{d}_3$ and transverse direction $\mathbf{d}_1$, this may be expressed as $\mathbf{f}_c = f_c\mathbf{d}_1$. The balance of linear momentum is then

$$\mathbf{n}'(s) = \rho g \mathbf{e}_y + f_c \mathbf{d}_1 \delta(s - s_c). \quad [45]$$

Here we have included self-weight with gravity $g$ acting in the negative $\mathbf{e}_y$ direction and linear density $\rho$. The delta function $\delta(s - s_c)$ accounts for contact at a single point, and creates a jump in the resultant force $\mathbf{n}$. Both $f_c$ and $s_c$ are unknown values that must be determined at each point in the evolution as part of the solution to the boundary value problem. To determine the two additional unknowns, the system requires two additional conditions, which are that the point $\mathbf{r}(s_c(t), t) = \mathbf{p}$, where $\mathbf{p}$ is the fixed contact point of the obstacle, and we highlight that the contact location along the rod may change with time. (Since the motion is restricted to a plane, this vector equation consists of the required two scalar conditions.)

In simulating this problem, we first integrate the system without contact, monitoring whether any point is near the obstacle, and stopping once a point along the rod first reaches the obstacle, i.e the first time $t = t^*$ at which there exists an $s = s^*$ for which $\mathbf{r}(s^*, t^*) = \mathbf{p}$. At this point, Eq. (45) has a solution with $f_c = 0, s_c = s^*$. For $t > t^*$, we then integrate the system with force balance (Eq. (45)). As a numerical shooting procedure, we integrate from $s = 0$ to $s = \ell$, in which case the other unknowns are the moment $\mathbf{m} = m\mathbf{e}_z$ and the force components $\mathbf{n} = n_x\mathbf{e}_x + n_y\mathbf{e}_y$ at $s = 0$. The 5 conditions to determine the shooting variables consist of the contact condition $\mathbf{r}(s_c(t), t) = \mathbf{p}$, and the three conditions that make up the free end boundary condition $\mathbf{m} = \mathbf{n} = \mathbf{0}$ at $s = \ell$. In this way, we employ standard continuation techniques to increment the system beyond $t^*$.

#### 7. Parameters and details of simulations

**A. Gravitropism: rotating base.** In simulating the rotating base under gravitropism (Fig. 4 main text), we orient the base at angle $\phi_0$ from the vertical $\mathbf{e}_z$ direction[‡]. Expressed in terms of the spherical unit vectors $\mathbf{e}_r = (\sin\phi_0, 0, \cos\phi_0)$, $\mathbf{e}_\phi = (\cos\phi_0, 0, -\sin\phi_0)$, $\mathbf{e}_\theta = (0,1,0)$, the frame at the point $S = 0$ is then given the form

$$\mathbf{d}_3(0,t) = \mathbf{e}_r \quad [46]$$

$$\mathbf{d}_1(0,t) = \cos(2\pi\omega t)\mathbf{e}_\phi + \sin(2\pi\omega t)\mathbf{e}_\theta \quad [47]$$

$$\mathbf{d}_2(0,t) = -\sin(2\pi\omega t)\mathbf{e}_\phi + \cos(2\pi\omega t)\mathbf{e}_\theta. \quad [48]$$

[‡] In terms of the angle $\theta$ appearing schematically in Fig. 3(A) of the main text, we have $\theta = \pi/2 - \phi_0$.

We fix $\omega = 1$, which is equivalent to scaling time based on the rotation rate of the base. We also set $\phi_0 = \pi/3$ and scale the total length $L = 1$. We simulate the gravitropic curvature laws with no axial growth and response rate $\mathcal{C}_{\text{grav}}$ taking values of $\mathcal{C}_{\text{grav}} = \{0.1, 1, 10, 50\}$. In this simulation we ignore the effect of self-weight, so that mechanical equilibrium is automatically satisfied with $\mathbf{u} = \hat{\mathbf{u}}$; we thus integrate Eqs. (12) and (13) to determine the morphology at each time step, and then update the curvature. Each parameter set is simulated up to time $t = 3$, which corresponds to three complete rotations of the base.

#### B. Phototropism.

**Fixed light source.** In simulating planar phototropism for a fixed light source (Fig. 5 of the main text), a light source is placed at the point $(1, 1)$, and the parameters $\ell = 1$, $U = 1$, and $\gamma \equiv 1$ (no axial growth) are fixed. This is equivalent to scaling time based on axial transport. The plant is clamped at the origin with tangent $\mathbf{d}_3 = (0, 1)$ at $s = 0$. We then simulate up to $t = 10$ for each combination of the parameter choices $Q = \{1, 5\}$, $\mathcal{C}_{\text{photo}} = \{0.5, 2.5\}$, to represent the different regimes of high and low uptake and phototropic response, respectively.

Note also that in simulating the time-delay differential equations, it is necessary to provide the form of the functions $\mathsf{e}_i$, $i = 1, 2$ for $-\ell/U \le t < 0$. These are chosen to be constant and equal to the value at $t = 0$, determined by the initial orientation.

**Moving light source - day/night cycle.** To simulate a day/night cycle (Fig. 6 of main text), we set $U = 1$, $\ell = 1$, $Q = 0.1$, $\gamma \equiv 1$, and $\mathcal{C}_{\text{photo}} = 1.5$. A light source with intensity $I(t) = \max\{0, \sin\omega t\}$ follows the path $\mathbf{p}(t) = (R\cos\omega t, Y, R\sin\omega t)$, where $\omega = 0.2$, $R = 3$, and $Y = 2$.

In the case of the additional autotropism terms, we increase $\mathcal{C}_{\text{photo}}$ to 3 and set $\xi = 0.3$. The increase in $\mathcal{C}_{\text{photo}}$ is chosen so that the motion during the day is similar to the non-autotropic case, as the autotropism serves to diminish the phototropic response in the presence of a stimulus. In both cases, one complete period is simulated, corresponding to day – when $I(t) > 0$, and night – when $I(t) = 0$.

#### C. Photogravitropism.

**Fixed light source.** For the simulations of main text Fig. 7 (A)-(E), we fix the parameters $U = 1$, $Q = 0.1$ and $\mathcal{C}_{\text{photo}} = 1$. Growth is uniform and linear: $\gamma = 1 + ct$ with $c = 0.1$, and initial length $L = 1$. A light source is placed at the point $\mathbf{p} = (4, 1)$. The plant is clamped at the origin with tangent $\mathbf{d}_3 = (0, 1)$ at $s = 0$. We then simulate up to $t = 10$ for each combination of the parameter choices $G = \{0.25, 2.5\}$, $\mathcal{C}_{\text{grav}} = \{0.1, 1\}$, . Here the parameter $G$ characterizes the effective impact of self-weight under gravity. In particular, by scaling rod length by $L$, moment by $E_b/L$ where $E_b$ is the bending stiffness, and noting that the gravitational force has magnitude $\rho g$, the non-dimensional moment balance equation, expressed in the reference variable $S$, is

$$m'(S) = -\gamma^2 G(S-1)\cos(\theta(S)), \quad G := \frac{\rho g L^3}{E_b} \qquad [49]$$

where $\theta$ is the angle between the tangent and the $x$-axis. In obtaining Eq. (49) we have used the geometric expression $\mathbf{r}'(S, t) = \gamma\mathbf{d}_3 = \gamma(\cos\theta\mathbf{e}_x + \sin\theta\mathbf{e}_y)$, and that the solution to the force balance $\mathbf{n}'(S) = \gamma\rho g\mathbf{e}_y$ subject to $\mathbf{n} = \mathbf{0}$ at $S = L$ is $\mathbf{n} = \rho g(S - L)\mathbf{e}_y$.

Thus, the parameter choices for $G$ and $\mathcal{C}_{\text{grav}}$ represent the different regimes of high and low mass/gravity and gravitropic response, respectively.

**Canopy escape.** In simulating the escape from shade (Fig. 7 (D)-(E) in main text), we have set $U = 1$, $Q = 0.5$, $\mathcal{C}_{\text{photo}} = 1$, $\mathcal{C}_{\text{grav}} = 0.1$, $\gamma = 1 + 0.25t$, $G = 0.05$ (see parameter description above). The initial length is $L = 1$, and the plant is clamped at the origin with tangent $\mathbf{d}_3 = (0, 1)$ at $s = 0$. The shade creating obstacle occupies the region $(x, y)$ with $x \le 1$, $y \ge 1.2$, so that the corner point and eventual contact point is $\mathbf{p} = (1, 1.2)$.

**D. Circumnutation.** For the simulations of circumnutation, main text Fig. 8 (A)-(B), the internal oscillator is located at the tip, with angular velocity $\omega = 1$; thus the period is $2\pi$ and we simulate one complete period. Plant length is scaled to $L = 1$, and axial growth is turned off ($\gamma \equiv 1$). In Fig. 8 (A) other parameters are $U = 5$, $Q = 5$, $\mathcal{C}_{\text{circ}} = 2$; in Fig. 8 (B) we use $U = 5$, $\mathcal{C}_{\text{circ}} = 1$, and we vary the uptake: $Q \in \{1, 2, 3\ldots, 10\}$. In Fig. 8 (C) the parameters are $U = 5$, $Q = 5$, $\mathcal{C}_{\text{circ}} = 1$, and the angular velocity is non-uniform; in particular the auxin gradient at the tip follows the line

$$\cos\theta x_1 + \sin\theta x_2,$$

with

$$\theta(t) = \omega t + \alpha\sin\hat{\omega}t.$$

The tip profiles in the figure are plotted for $\omega = 1$, $\hat{\omega} = 5$, and varying $\alpha = \{0, 0.15, 0.3, \ldots, 1.5\}$.

In these simulations we have also given the plant an initial curvature, which serves to better center the motion about the base of the plant, for visualization purposes (the initial curvature only creates a translation of the tip pattern). The initial curvatures used were as follows: $\mathsf{u}_1 = 0$ in Fig. 8 (A)-(C), while $\mathsf{u}_2 = 1.25$ in Fig. 8 (A), $\mathsf{u}_2 = 0.5$ in Fig. 8 (B), and $\mathsf{u}_2 = 0.45 - 0.07\alpha$ in Fig. 8 (C) (this choice was made to avoid overlapping of the tip patterns with varying $\alpha$).

**E. Thigmotropism.** In simulating thigmotropism, main text Fig. 8 (D)-(F), we have set $\mathcal{C}_{\text{thig}} = 10$, and varied the uptake $Q$ and angular velocity $\omega$ as follows: $Q = 3, \omega = 2$ in Fig. 8 (D), $Q = 3, \omega = 6$ in Fig. 8 (E), and $Q = 5, \omega = 6$ in Fig. 8 (F). Again, axial growth is turned off and the plant length is $L = 1$. In each case total simulation time is $t = 10$. In the thigmotropism formulation, with the signal coming from a single point, the curvatures may be determined exactly, given by

$$u_1 = -\mathcal{C}_{\text{thig}} \exp\left(-\frac{QS}{U}\right) \sin\left(\psi_0 + \frac{\omega S}{U}\right) t \tag{50}$$

$$u_2 = \mathcal{C}_{\text{thig}} \exp\left(-\frac{QS}{U}\right) \cos\left(\psi_0 + \frac{\omega S}{U}\right) t. \tag{51}$$

Here the angle $\psi_0$ indicates the point of contact (which is set at $s = 0$). In the presented simulations, $\psi_0 = \pi/2$, so that the contact point is at $\mathbf{r}(0, t) + a\mathbf{d}_2$, where $a$ is the cross-sectional radius, which was fixed at $a = 0.02$. From the formulas in Section 3, we then obtain that the curvature $\kappa$ and torsion $\tau$ will evolve according to

$$\kappa = \mathcal{C}_{\text{thig}} \exp\left(-\frac{QS}{U}\right) t \tag{52}$$

$$\tau = \frac{\omega}{U}. \tag{53}$$

Note that a helix of radius $\alpha$ and pitch $\beta$ (i.e. where the angle of the helix $\phi$ satisfies $\tan\phi = \beta/\alpha$) has curvature $\hat{\kappa} = \alpha/\left(\alpha^2 + \beta^2\right)$ and torsion $\hat{\tau} = \beta/\left(\alpha^2 + \beta^2\right)$. Since the torsion of the plant is fixed by the ratio of rotational to axial auxin velocity, Eq. (53), and the helical radius for a pole of radius $c$ and plant radius $a$ is $\alpha = c + a$, we can solve for the pitch, or equivalently the angle $\phi$, which satisfies $\sin(2\phi) = \omega(a + c)/U$. It follows that the curvature $\hat{\kappa} = \cos^2\phi/(a + c)$; in our simulations we have fixed the pole radius $c = 0.05$. In this formulation, the curvature increases linearly in time at every point. This unrealistic (in long times) aspect could be corrected by having a depleting auxin source at the contact point. However, in any case we must account for the fact that the curvature cannot increase beyond $\hat{\kappa}$, simply due to the presence of the pole. Thus, in simulating the wrapping around a pole, at each spatial point we increase the curvatures according to Eqs. (50) and (51), until the curvature $\kappa = \hat{\kappa}$, where $\kappa$ is given by Eq. (52), at which point we freeze the curvatures in the simulation (in this way, we account for the fact that the intrinsic curvature may keep increasing, but the actual curvature may not due to the mechanical contact, while avoiding the problem of having to compute the mechanical contact force density).

**F. Pole dance.** In SI movies, we include a simulation that consists of a plant that searches for a pole via the circumnutation model, while also undergoing axial growth, and then begins to wrap around it following the thigmotropism model once contact is made. In this simulation, the parameters used were $U = 6$, $Q = 10$, $\mathcal{C}_{\text{circ}} = 3$, and circumnutation oscillator frequency $\omega = 5$ originating at the base $S = 0$. The plant is clamped at the origin, has radius $a = 0.025$, is initially straight and has initial length $L = 1$ and growth rate $\frac{\partial\gamma}{\partial t} = 0.4$. A vertical pole with radius $c = 0.05$ passes through the point $\{-0.68, -0.52, 0\}$. The plant first makes contact with the pole at time $t = 3$, and at contact point defined by reference arc length $S_c = 0.8$ and angle $\psi_c = 2.26$[§].

Once contact is made, we turn off the circumnutation signal, and only evolve the portion of the plant, $S > S_c$, i.e. from pole to end. This follows the thigmotropism curvature evolution, with parameters $U = 1$, $Q = 3$, $\mathcal{C}_{\text{thig}} = 9$, and $\omega = 0.77$. The choice of $\omega$ is made for computational convenience, as this particular value means that the pitch of the helix is exactly equal to the angle at which contact is made, and no rotation of the tangent about the contact point is needed. The wrapping portion of the evolution is simulated from $t = 3$ up to $t = 4.5$.

#### 8. Description of Movies

**SI movie S1:** Gravitropism with rotating base, and gravitropic response parameter $C_{\text{thig}} = 0.1$. Other simulation parameters provided in SI Section 7.

**SI movie S2:** Gravitropism with rotating base, and gravitropic response parameter $C_{\text{thig}} = 1$. Other simulation parameters provided in SI Section 7.

**SI movie S3:** Gravitropism with rotating base, and gravitropic response parameter $C_{\text{thig}} = 10$. Other simulation parameters provided in SI Section 7.

**SI movie S4:** Gravitropism with rotating base, and gravitropic response parameter $C_{\text{thig}} = 50$. Other simulation parameters provided in SI Section 7.

**SI movie S5:** Phototropism, simulation of a day-night cycle, with no autotropism. Simulation parameters provided in SI Section 7.

[§] Rather than define the location of the pole, we have defined the location and time of the contact, and used these to define the pole; we then verify that with the pole defined in this way, no prior contact was made.

**SI movie S6:** Phototropism, simulation of a day-night cycle, with autotropism. Simulation parameters provided in SI Section 7.

**SI movie S7:** Thigmotropism, pole wrapping, with low uptake ($Q = 3$) and low angular velocity ($\omega = 2$). Other simulation details provided in SI Section 7.

**SI movie S8:** Thigmotropism, pole wrapping, with low uptake ($Q = 3$) and high angular velocity ($\omega = 6$). Other simulation details provided in SI Section 7.

**SI movie S9:** Thigmotropism, pole wrapping, with high uptake ($Q = 5$) and low angular velocity ($\omega = 6$). Other simulation details provided in SI Section 7.

**SI movie S10:** Pole dance. Circumnutation with axial growth, followed by thigmotropic pole wrapping. Simulation parameters provided in SI Section 7.


1. Moulton DE, Lessinnes T, Goriely A (2020) Morphoelastic rods III: Differential growth and curvature generation in elastic filaments.
2. Band LR, et al. (2014) Systems analysis of auxin transport in the Arabidopsis root apex. *The Plant Cell* 26(3):862–875.
3. Fozard J, Bennett M, King J, Jensen O (2016) Hybrid vertex-midline modelling of elongated plant organs. *Interface Focus* 6(5):20160043–14.
4. Chauvet H, Moulia B, Legué V, Forterre Y, Pouliquen O (2019) Revealing the hierarchy of processes and time-scales that control the tropic response of shoots to gravi-stimulations. *Journal of experimental botany* 70(6):1955–1967.
5. Band LR, King JR (2011) Multiscale modelling of auxin transport in the plant-root elongation zone. *Journal of mathematical biology* 65(4):743–785.
6. Silk WK (1989) On the Curving and Twining of Stems. *Environmental and Experimental Botany* 29(1):95–109.
7. Reinhold L, Sachs T, Vislovska L (1972) The role of auxin in thigmotropism in *Plant Growth Substances 1970.* (Springer), pp. 731–737.
8. Shabala SN, Newman IA (1997) Root nutation modelled by two ion flux-linked growth waves around the root. *Physiologia Plantarum* 101:770–776.
9. Stolarz M (2014) Circumnutation as a visible plant action and reaction. *Plant Signaling & Behavior* 4(5):380–387.
10. Galland P (2002) Tropisms of avena coleoptiles: sine law for gravitropism, exponential law for photogravitropic equilibrium. *Planta* 215(5):779–784.
