## Supplementary material for "Multiscale integration of environmental stimuli in plant tropism produces complex behaviors": Videos: SM movie captions.rtf

SM movie 1: Gravitropism with rotating base, and gravitropic response parameter $C_\text{thig}=0.1$.Other simulation parameters provided in SM Section 7.

SM movie 2: Gravitropism with rotating base, and gravitropic response parameter $C_\text{thig}=1$. Other simulation parameters provided in SM Section 7.

SM movie 3: Gravitropism with rotating base, and gravitropic response parameter $C_\text{thig}=10$. Other simulation parameters provided in SM Section 7.

SM movie 4: Gravitropism with rotating base, and gravitropic response parameter $C_\text{thig}=50$. Other simulation parameters provided in SM Section 7.

SM movie 5: Phototropism, simulation of a day-night cycle, with no autotropism. Simulation parameters provided in SM Section 7.

SM movie 6: Phototropism, simulation of a day-night cycle, with autotropism. Simulation parameters provided in SM Section 7.

SM movie 7: Thigmotropism, pole wrapping, with low uptake ($Q=3$) and low angular velocity ($\omega=2$). Other simulation details provided in SM Section 7. 

SM movie 8: Thigmotropism, pole wrapping, with low uptake ($Q=3$) and high angular velocity ($\omega=6$). Other simulation details provided in SM Section 7. 

SM movie 9: Thigmotropism, pole wrapping, with high uptake ($Q=5$) and low angular velocity ($\omega=6$). Other simulation details provided in SM Section 7. 

SM movie 10: Pole dance. Circumnutation with axial growth, followed by thigmotropic pole wrapping. Simulation parameters provided in SM Section 7.
