## Supplementary figures and images for "Multiscale integration of environmental stimuli in plant tropism produces complex behaviors"

### SI_movie_S1.gif

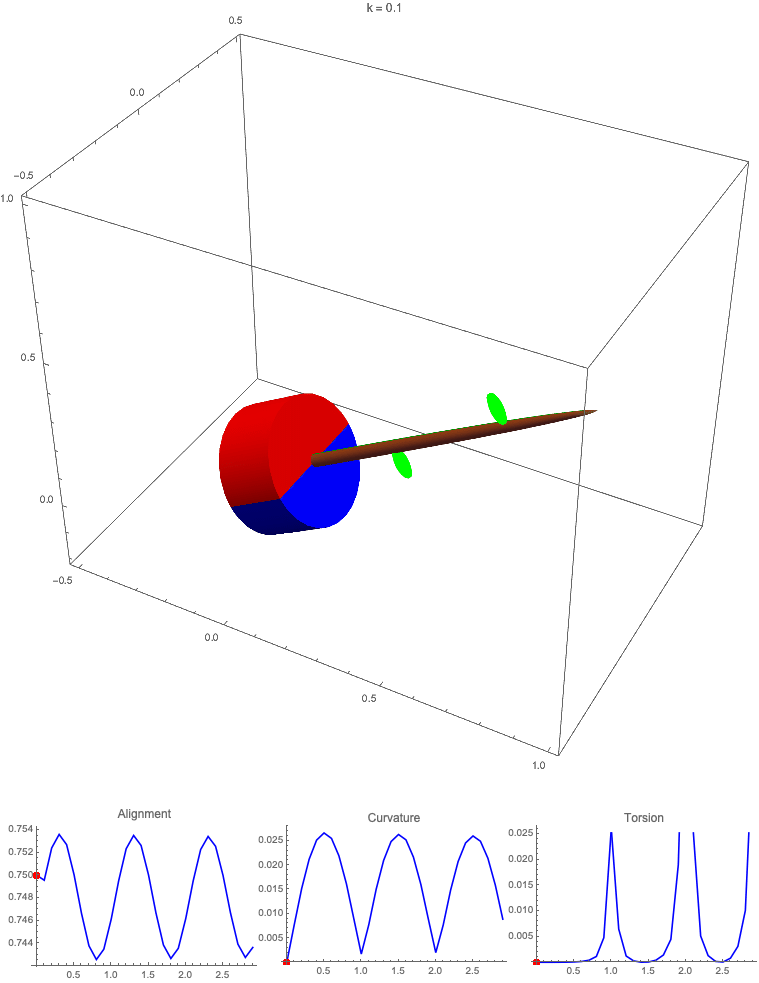

### SI_movie_S2.gif

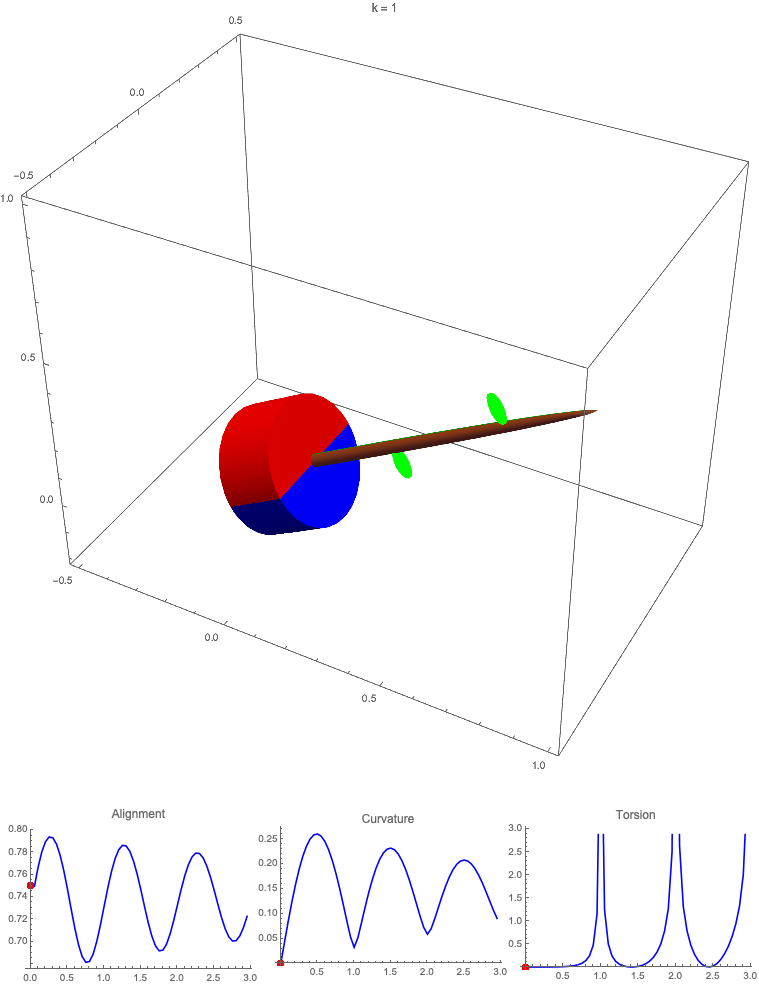

### SI_movie_S3.gif

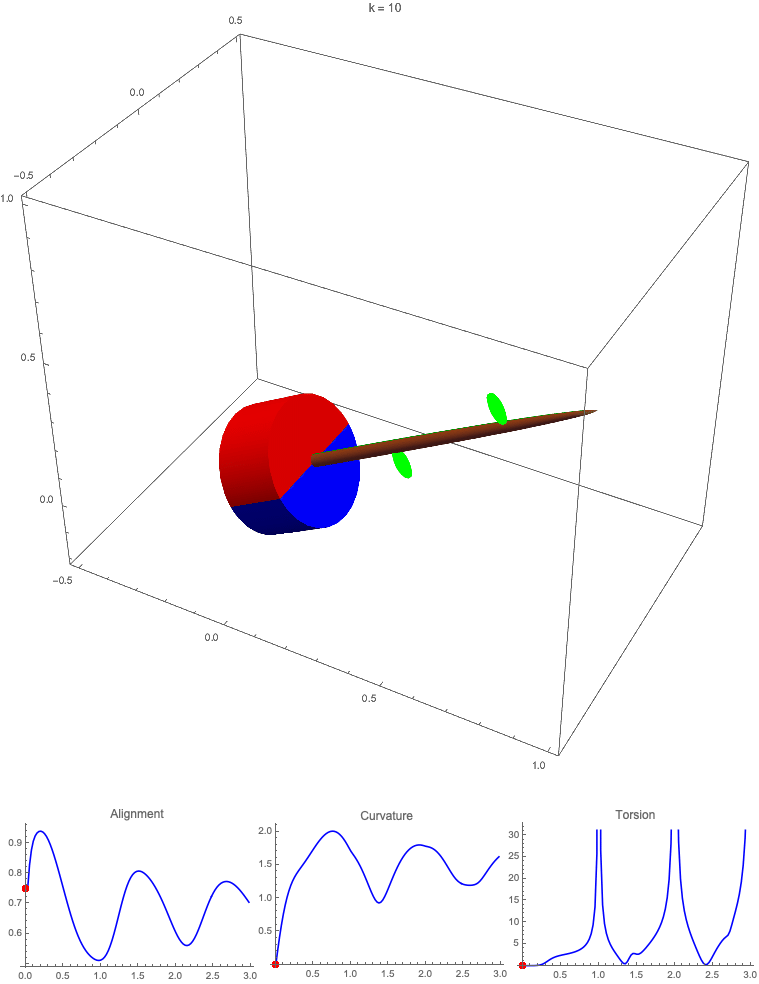

### SI_movie_S4.gif

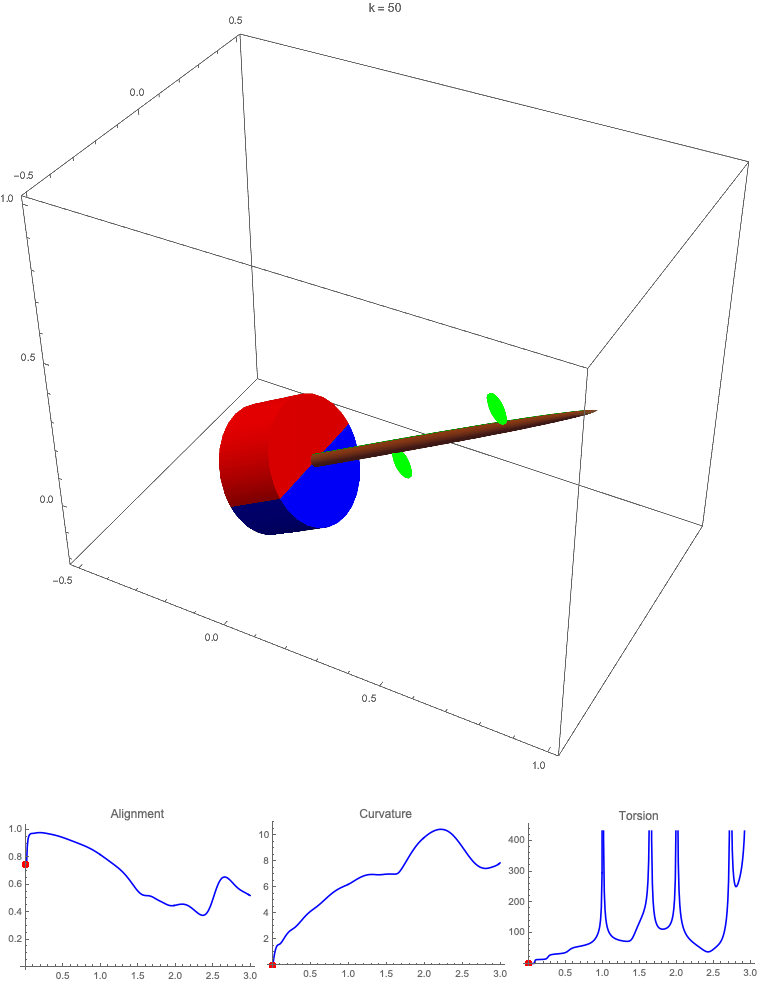

### SI_movie_S5.gif

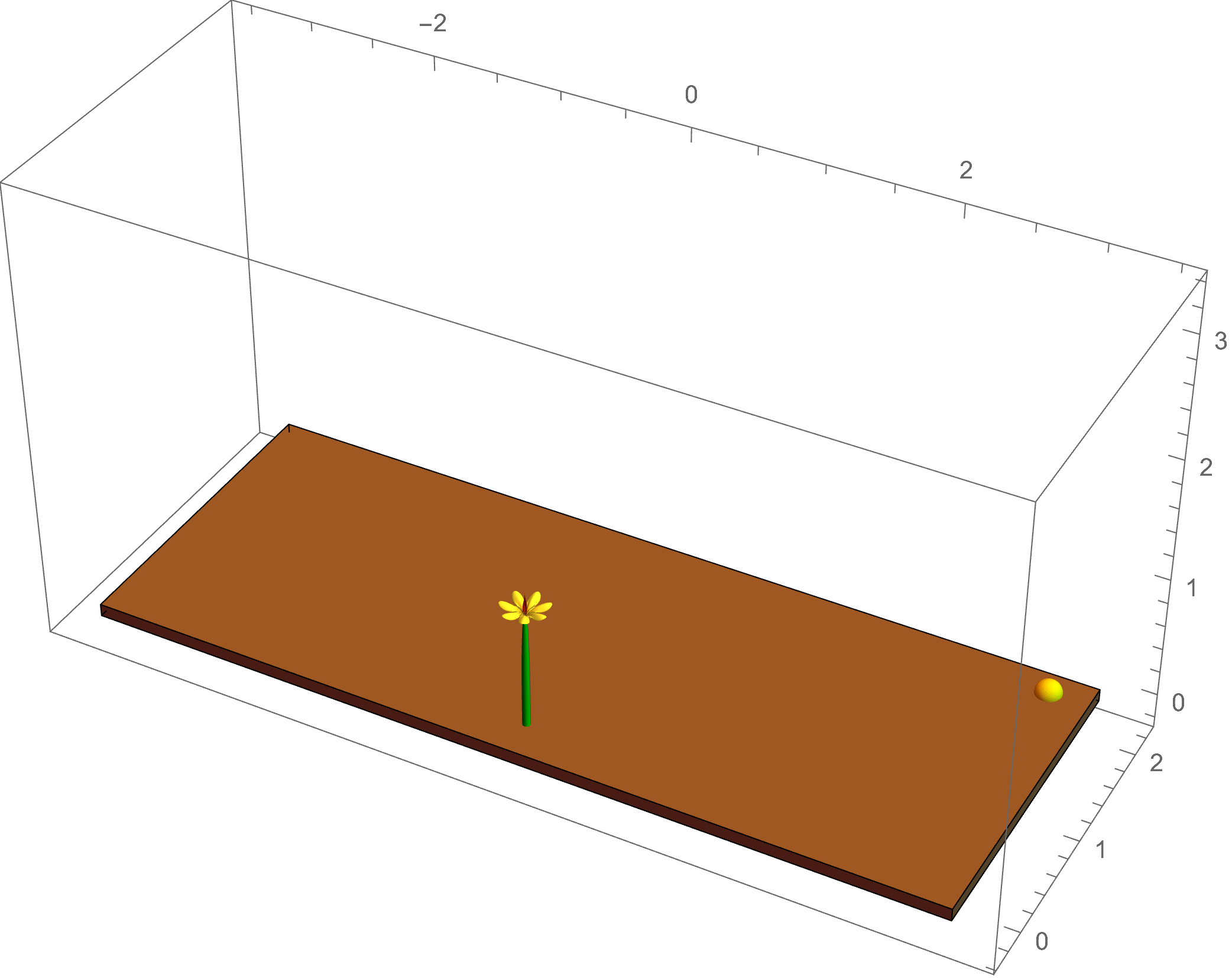

### SI_movie_S6.gif

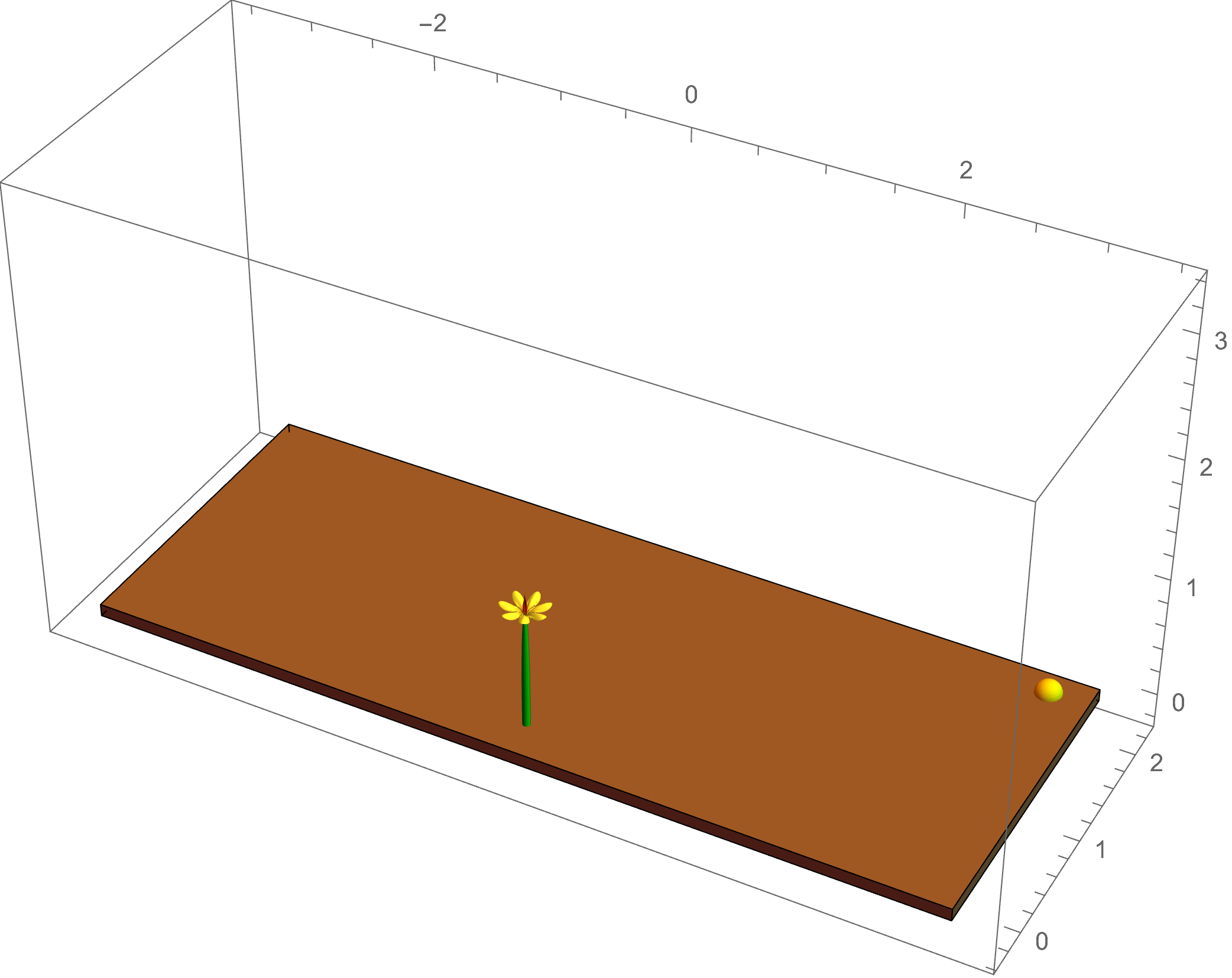

### SI_movie_S7.gif

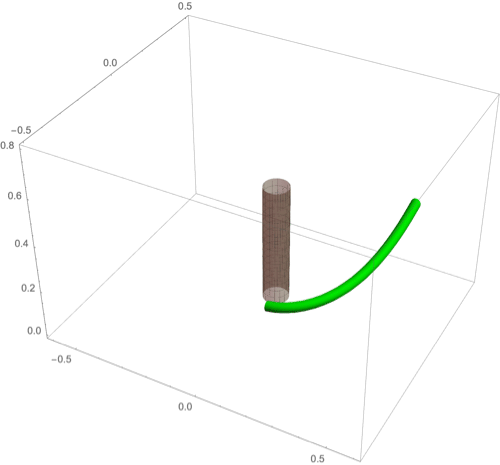

### SI_movie_S8.gif

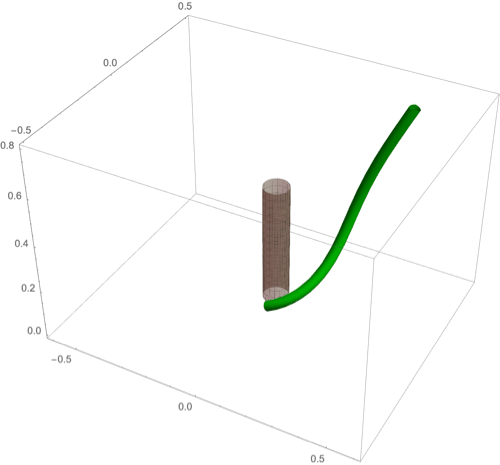

### SI_movie_S9.gif

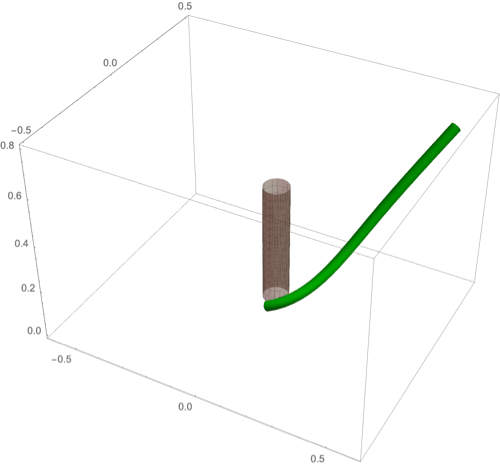

### SI_movie_S10.gif

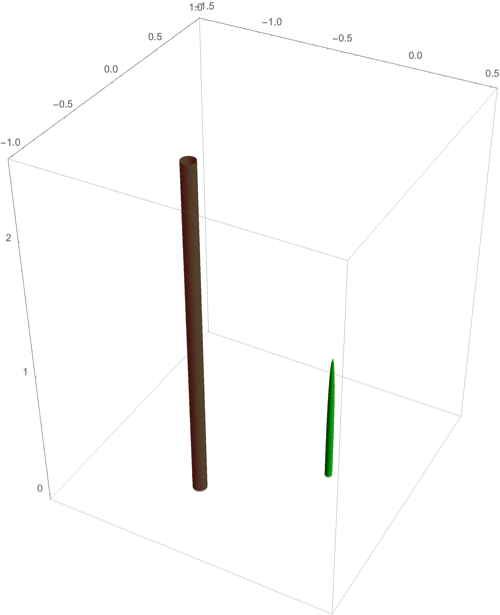
